# Cholesterol availability drives the assembly of mitochondria-associated ER membranes

**DOI:** 10.64898/2026.10.29.762740

**Authors:** M Uceda, K Kabra, A.C Ferre, L Serrano-Cuñarro, N Gomez-Lopez, M Pera, D Larrea, R.R Agrawal, K.A Tamucci, C Anton-Barros, T.D Yun, K.R Velasco, E.A Schon, J Montesinos, E Area-Gomez

**Author notes:** Co-corresponding authors. Please address correspondence to;.

## Abstract

Mitochondria-associated endoplasmic reticulum (ER) membranes (MAM domains, or MAMs) are transient lipid raft domains of the ER that coordinate multiple cellular metabolic pathways and the ER-mitochondria communication. However, the mechanisms that drive their formation remain poorly understood. Here, we reveal that MAM assembly is dynamically regulated by the availability of plasma membrane (PM) cholesterol. We show that cholesterol delivered through high-density lipoproteins (HDLs) via the scavenger receptor class B type 1 (SR-B1) pathway is rapidly redistributed from the PM to the ER, where increased cholesterol availability promotes the assembly of MAM domains. In turn, MAM formation promotes the communication between the ER and mitochondria, the esterification of excess cholesterol and the suppression of cholesterol biosynthesis. Disruption of MAMs uncouples these responses, altering cholesterol trafficking and metabolic adaptation to extracellular cholesterol. Our findings identify MAM as a dynamic metabolic regulator that translates changes in extracellular lipoprotein availability into coordinated intracellular metabolic responses and cellular adaptation.

## INTRODUCTION

The establishment of functional contact sites between organelles has fundamentally changed our understanding of cellular regulation. Nevertheless, important questions remain regarding how these contacts are formed and dynamically regulated, and how they contribute to coordinate the multiple pathways that govern cellular metabolism [1].

Among the best-characterized contact sites are the ones between mitochondria and the endoplasmic reticulum (ER), also known as mitochondria-ER contact sites, or MERCs [2]. The formation of MERCs is associated with the establishment of transient lipid membrane domains within the ER, known as mitochondria-associated ER membranes (MAM), through mechanisms that remain incompletely understood. MAM domains in the ER transiently cluster and compartmentalize enzymes and regulatory factors involved in multiple metabolic pathways, including calcium homeostasis, mitochondrial dynamics, and lipid metabolism – in particular, cholesterol homeostasis [2, 3]. However, the molecular mechanisms that drive the formation, remodeling, and regulation of these specialized ER domains remain largely unknown.

Cholesterol is essential for the regulation of the structure and function of biological membranes. Due to its planar conformation, cholesterol can fit into lipid bilayers and modify membrane properties such as thickness, fluidity, permeability, and curvature [4]. Moreover, cholesterol levels above a certain threshold cause membrane partitioning and the formation of lipid raft (LR) domains [5, 6]. These transient liquid-ordered (Lo) domains, which are typically surrounded by and segregated from the more abundant liquid-disordered (Ld) domains, carry specific subsets of lipid-binding proteins to facilitate conformational changes and/or protein interactions for the regulation of cell signaling, trafficking, and protein sorting [7]. Therefore, cholesterol modulates membrane properties in a concentration-dependent manner, and its distribution can be considered a signaling mechanism, involved in vesicle formation, membrane fusion, and protein sorting, with distinct effects across different cellular and membrane systems [8, 9].

At the molecular level, the cell employs many highly regulated mechanisms to sense and maintain cholesterol levels in each cellular and intracellular membrane [9]. The plasma membrane (PM) contains the highest levels of cholesterol, divided into pools with different accessibility to enzymatic activity. Beyond a threshold, excess cholesterol in the PM becomes accessible and is transported to the ER. Despite being only a small fraction of total cellular cholesterol, this ER pool acts as a central cholesterol sensor, repressing cholesterol biosynthesis at the transcriptional and post-transcriptional levels, while inducing its esterification to maintain homeostasis [10]. However, the identity, localization, and regulation of these pools of cholesterol in the ER are not completely understood.

To fulfill their cholesterol needs, most cell types rely on the uptake of low-density lipoproteins (LDLs), very low-density lipoproteins (VLDLs) and high-density lipoproteins (HDLs) [11, 12]. In most human cells, extracellular cholesterol is primarily internalized from LDL particles, which are endocytosed and fuse with lysosomes, triggering the hydrolysis of their cholesterol esters (CE) into free cholesterol [13]. To a lesser degree, human cells can also acquire cholesterol from the extracellular medium by the non-vesicular internalization of HDLs via Aster proteins [10, 14], although this type of lipoprotein is considered to be mainly involved in cholesterol efflux [15, 16]. However, published evidence indicates that cells can internalize these particles through HDL-avid receptors, such as the scavenger receptor B1 (SR-B1; gene *Scarb1*), to provide cells with extra cholesterol under specific conditions, such as steroidogenesis or LDL-deficiencies [17]. Furthermore, recent mechanistic and translational studies indicate that HDLs can elicit multiple cellular responses, different from those of LDL and VLDL, such as enhancing glucose uptake and mitochondrial activity, as well as having antioxidative and anti-apoptotic properties [18, 19], but the molecular mechanism behind these effects remain unclear.

Here, we demonstrate that the mobilization of excess cholesterol from the PM to the ER triggers the formation of MAM domains and the functional interaction between ER and mitochondria, as well as the inhibition of cholesterol biosynthesis. Moreover, we show that cholesterol internalization and MAM formation are specifically stimulated by the uptake of HDLs, compared to other lipoproteins. Taken together, our data suggest that MAMs are cholesterol-rich regulatory pools in the ER, and provide a new framework understanding how different extracellular lipoproteins modulate intracellular metabolic feedback responses.

## RESULTS

### Cholesterol internalization and its delivery to the ER induces the formation of MAM domains

Previous work suggests that, as a lipid raft-like domain, the formation of MAM is triggered by elevations in cholesterol levels in the ER, likely delivered from the PM [20]. To validate these data, we quantified MAM activity after stimulating cholesterol uptake and traffic from the PM to the ER by incubating cultured cells with increasing concentrations of exogenously-added cholesterol, as shown before [21].

Specifically, we incubated mouse embryonic fibroblasts (MEFs) and human neuroblastoma cells (SH-SY5Y) in media containing 3% serum supplemented with increasing concentrations of water-soluble cholesterol (0.5 mg, 3 mg, or 5 mg per 100 mL) spiked with ^3^H-cholesterol, and measured its internalization and esterification over time. Under these conditions, cultured cells displayed a dose-dependent increase in cholesterol uptake (**Figs. 1A and S1A)**, resulting in the stimulation of MAM functions, including the esterification of cholesterol by acyl-CoA:cholesterol acyltransferase 1 (ACAT1, gene *Soat1*) [22] (**Figs. 1B and S1B)** and the synthesis and transfer of phosphatidylserine (PtdSer) to mitochondria for its decarboxylation into phosphatidylethanolamine (PtdEtn) (**Figs. 1C and S1C**) [23]. Likewise, incubation with fluorescently-labeled cholesterol (NBD-cholesterol) showed a time-dependent uptake (**Fig. 1D**) and induced an increase in MAM function (**Fig. 1E**), as well as a significant redistribution of endogenous cholesterol towards perinuclear areas over time (**Figs. 1F-H**). Conversely, inhibition of 3-hydroxy-3-methylglutaryl-CoA reductase (HMGCR), the first and rate-limiting enzyme in the mevalonate pathway [11], by atorvastatin did not significantly impact MAM activities (**Fig. 1I**), indicating that the formation of this domain is not induced by *de novo* synthesized cholesterol.

**Fig. 1.**
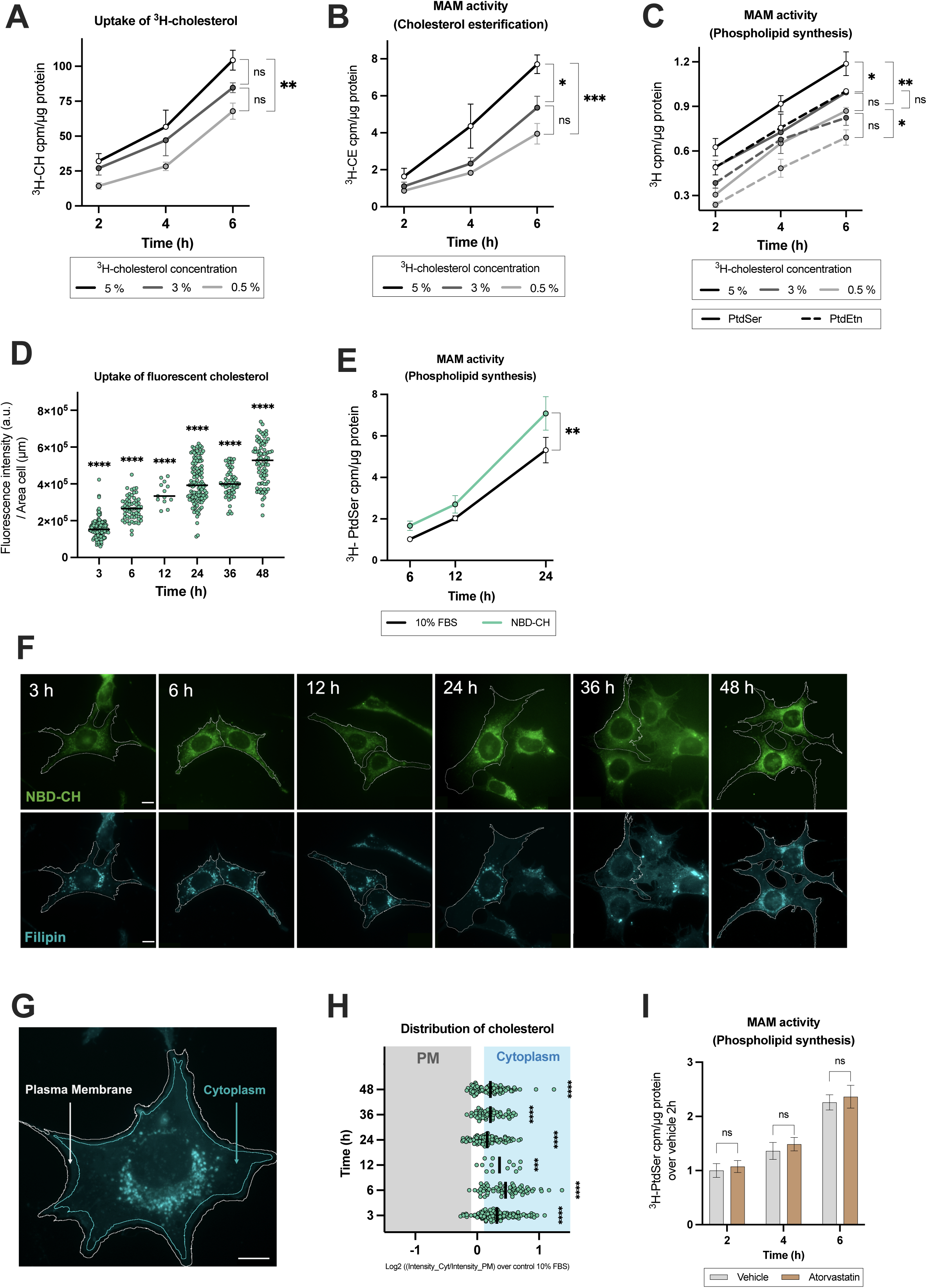
Cholesterol internalization induces MAM formation. **(A)** Quantification of exogenous ^3^H-cholesterol incorporation to MEF cells at increasing concentrations and times of incubation (*n = 3 biological replicates*). **(B-C)** Analysis of MAM activity in MEF cells exposed to increasing concentrations of exogenous ^3^H-cholesterol, quantified by **(B)** cholesterol esterification (CE) and (**C**) the incorporation of ^3^H-serine into phosphatidylserine (PtdSer) and phosphatidylethanolamine (PtdEtn) (*n = 3 biological replicates*). **(D)** Quantification of fluorescently-labeled cholesterol internalization in MEF cells at the indicated times (*n = 59-122 cells per condition)*. **(E)** Quantification of MAM activity in MEF cells exposed to NBD-cholesterol over time, measured by PtdSer synthesis (*n = 4 biological replicates*). **(F)** Representative images of cells exposed to NBD-cholesterol signal and stained with filipin at the indicated incubation times. Scale bar = 10 μm. **(G)** Representative image of cells stained with filipin and the distribution of cholesterol in plasma membrane and cytoplasm areas. Scale bar = 10 μm. **(H)** Quantification of the distribution of cholesterol in MEF cells incubated with NBD-cholesterol at the indicated times. (*n = 59-122 cells per condition)*. **(I)** Measurement of MAM activity in MEF cells pre-treated with atorvastatin (20 μM) for 16h, measured by PtdSer synthesis (*n = 3 biological replicates*). All data are expressed as the mean ± SEM, * p<0.05, ** p<0.01, *** p<0.001, **** p<0.0001.

To validate this result, we induced the mobilization of cholesterol in the PM towards the ER by adding exogenous sphingomyelinase (SMase) to cell cultures [24] which, after sphingomyelin hydrolysis, makes cholesterol accessible to be trafficked to the ER. As expected, SMase treatment induced the internalization of cholesterol, as measured by the uptake of ^3^H-labeled cholesterol and its delivery to the ER (**Fig. 2A**), thereby inducing the redistribution of cholesterol towards perinuclear areas, as shown by filipin staining (**Fig. 2B**). Concomitantly, this stimulation of cholesterol trafficking to the ER resulted in an increase in ER-mitochondria apposition or contacts (MERCs) (**Fig. 2C**), as well as the activation of MAM functions, including cholesterol esterification by ACAT1 (**Figs. 2D and S2A)** and phospholipid synthesis (**Figs. 2E and S2B)**. In agreement, MAM activation was attenuated by both the inhibition of endogenous SMase by GW4869 [20] (**Fig S2B**) and the inhibition of cholesterol trafficking by U18666A (**Fig. 2E**) and by AI-d3, an inhibitor of Aster proteins [14] ( **Fig. S2C**).

**Fig. 2.**
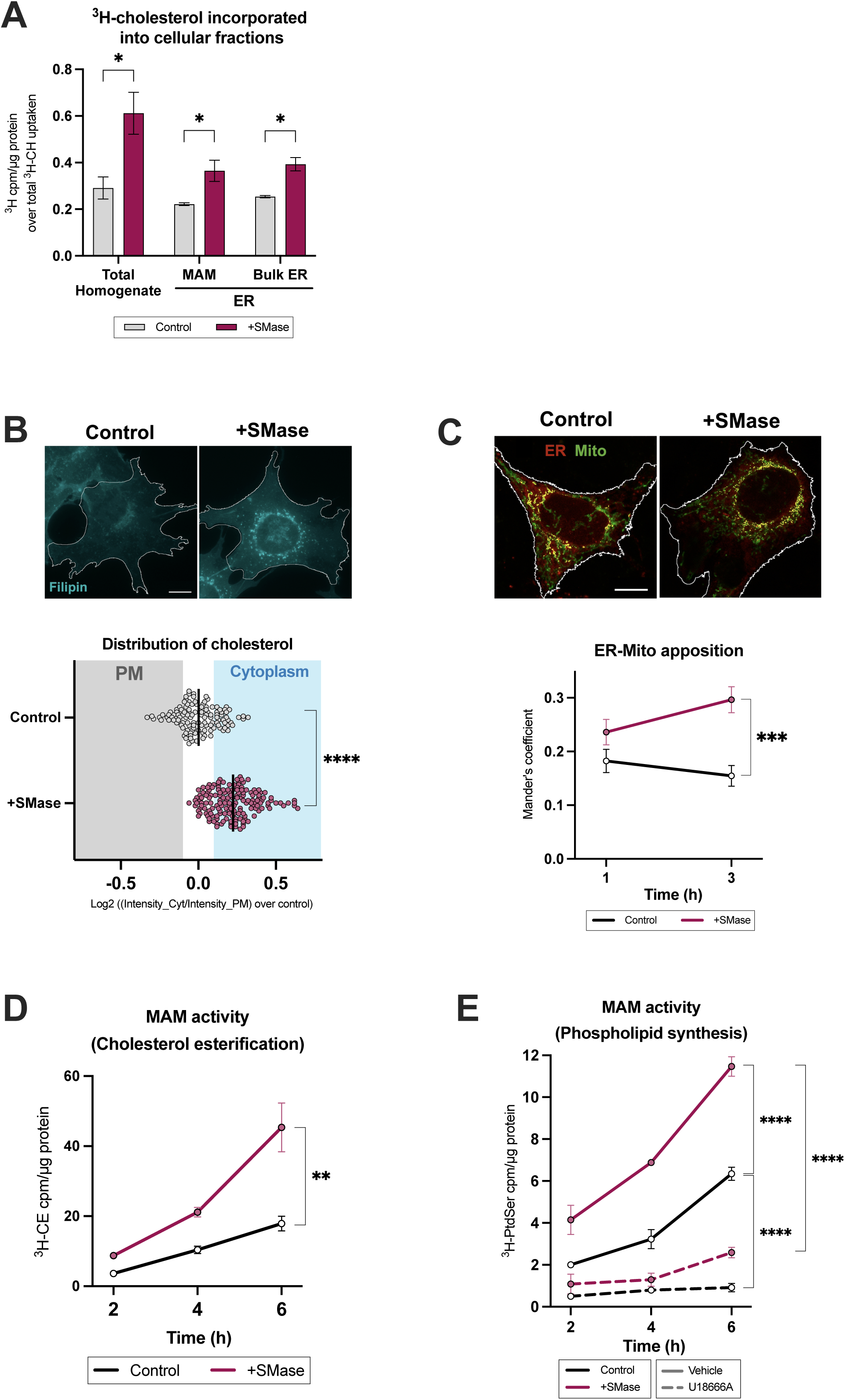
Cholesterol delivery to the ER induces MAM formation. All experiments were performed in MEF cells exposed to 1U/mL SMase in 5% FBS media for the indicated times. **(A)** Quantification of ^3^H-cholesterol incorporation into total homogenate and different ER fractions (bulk ER, MAM) (*n = 3 biological replicates*). **(B)** Representative images of cholesterol distribution by filipin staining and the quantification of fluorescent signal (lower panel) in cells treated with SMase for 3h. Quantification below.. Scale bar = 10 μm (*n = 121-162 cells per condition*). **(C)** Representative image of ER (red) and mitochondria (green) apposition after SMase treatment for the indicated times. Quantification below (*n = 43 cells per condition*) Scale bar = 10 μm. **(D)** Measurement of MAM activity, assessed by cholesterol esterification into cholesteryl esters (*n = 3 biological replicates*). **(E)** Measurement of MAM activity in cells exposed to 1 μM U1866A over the indicated times, as measured by phosphatidylserine (PtdSer) synthesis (*n = 3 biological replicates*). All data are expressed as the mean ± SEM, * p<0.05, ** p<0.01, *** p<0.001, **** p<0.0001.

Our data thus indicate that cholesterol mobilization from the PM, rather than its biosynthesis, induces transient increases in cholesterol in the ER and the consequent formation and activation of MAM domains, and thus contact of the ER with mitochondria.

Cholesterol delivery to the ER represses cholesterol biosynthesis by inhibiting the activity of specific rate-limiting enzymes and repressing SREBP2 (gene *Srebf2*) processing and expression. In agreement, our data show that SMase treatment and subsequent MAM activation resulted in reduced expression and activity of the cholesterol biosynthetic rate-limiting enzymes, such as HMGCR (**Figs. 3A and 3B**). SMase treatment did not affect the transcription levels of other enzymes involved in cholesterol biosynthesis, such as squalene synthase (SQS; gene *Fdft1*), squalene monooxygenase epoxidase (SQLE, gene *Sqle*), or lanosterol synthase (LSS, gene *Lss*) (**Fig. S3A**).

**Fig. 3.**
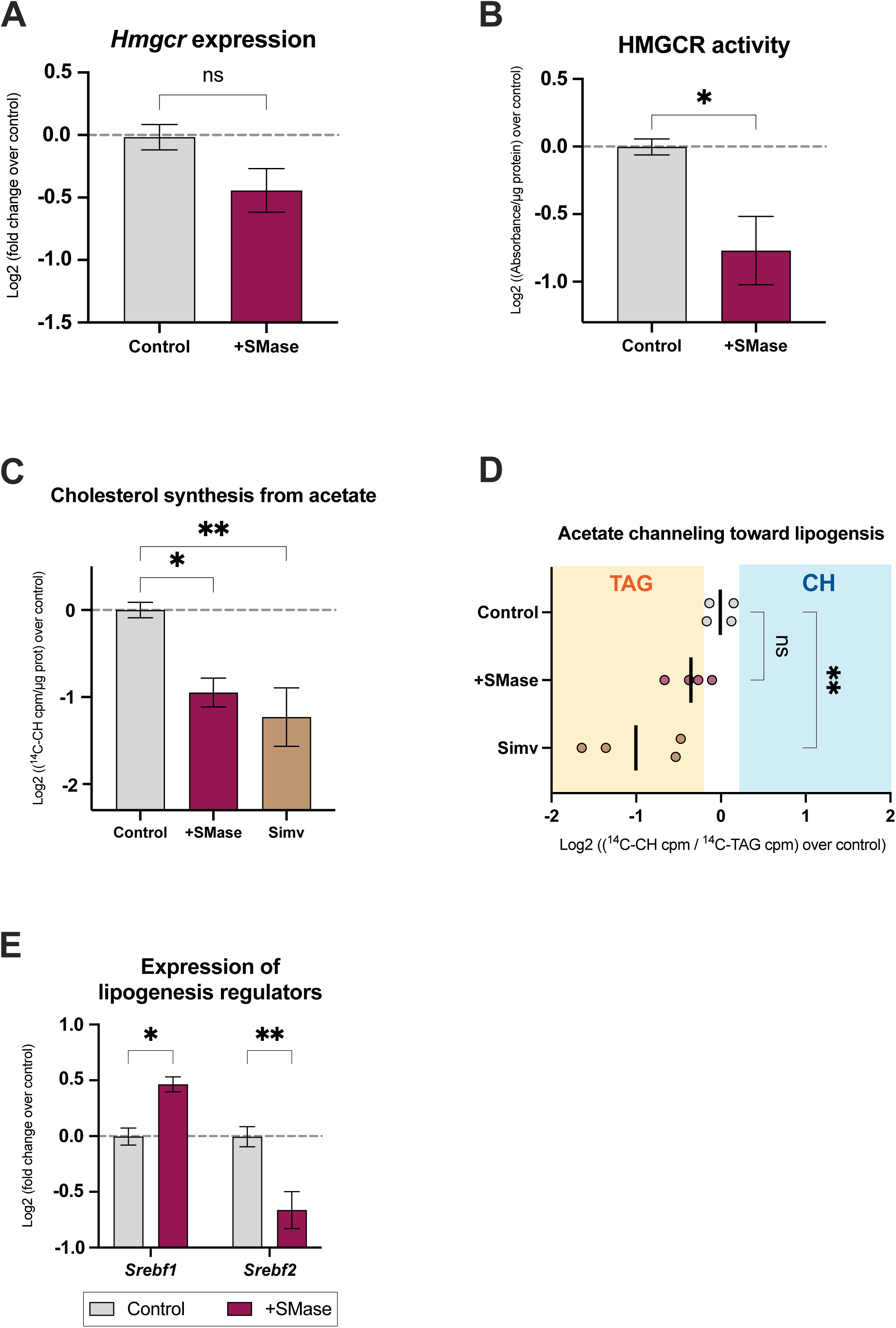
Cholesterol delivery to the ER represses cholesterol biosynthesis. All experiments were performed in MEF cells exposed to 1U/mL SMase in 5% FBS media for the indicated times. **(A)** Measurement of mRNA expression levels of *Hmgcr* (*n = 3 biological replicates*). **(B)** Measurement of HMGCR activity (*n = 3 biological replicates*). **(C)** Quantification of *de novo* cholesterol synthesis from ^14^C-acetate in cells exposed to 1 U/mL SMase or simvastatin (Simv) (*n = 4 biological replicates*). **(D)** Analysis of the incorporation of ^14^C-acetate into *de novo* synthesized cholesterol (CH) or triglycerides (TAG) in cells exposed to 1U/mL SMase or simvastatin (Simv) (*n = 4 biological replicates*). **(E)** Quantification of mRNA levels of *Srebf1* and *Srebf2* (*n = 3 independent experiments*). All data are expressed as the mean ± SEM, * p<0.05, ** p<0.01.

To determine the effect of MAM formation on cholesterol biosynthesis, we incubated cells with ^14^C-acetate, the primary precursor of this pathway, and measured its incorporation into newly-synthesized cholesterol at 3 hours in 5% FBS media, in the presence or absence of SMase treatment. Consistent with previous reports [24], SMase treatment resulted in a reduction in cholesterol biosynthesis, similar to that shown by cells treated with simvastatin, an HMGCR inhibitor (**Fig. 3C**).

Concomitantly, SMase and statin-treated cells presented with increases in newly synthesized triglycerides (TAGs), suggesting a shift in the use of acetate toward glycolipid production (**Fig. 3D**). This shift was accompanied by reductions in the expression of *Srebf2*, the master regulator of cholesterol biosynthesis, and increases in *Srebf1*, which is mainly involved in fatty acid and TAG synthesis [25] (**Figs. 3E and S3B**).

Altogether, these findings indicate that the mobilization of cholesterol from the PM to the ER induces a feedback response that suppresses cholesterol biosynthesis while redirecting acetate-dependent lipogenesis toward TAG synthesis. This trafficking of cholesterol correlates with MAM formation, suggesting that the contact site between the ER and mitochondria might be involved in lipogenesis balance.

### MAM formation induces cholesterol internalization and its delivery to the ER

To further test the relevance of MAM regulation in the control of cholesterol metabolism, we next measured cholesterol uptake and synthesis in cell models where MAM activities are impaired or stimulated due to the ablation of mitofusin 2 (MFN2-KO cells) or presenilins −1 and −2 (PSEN1/2-KO; PS-DKO cells), respectively, as previously shown [26] **(Fig. S4A)**

Consistent with MAM activity in these models, PS-DKO cells displayed a redistribution of endogenous cholesterol toward a more perinuclear localization, suggesting enhanced cholesterol mobilization from the PM (**Fig. 4A**). In contrast, MAM-deficient MFN2-KO cells showed the opposite phenotype (**Fig. 4A**). As previously shown [20], MAM upregulation in PS-DKO cells is associated with increased uptake of extracellular cholesterol, similar to cells treated with SMase and contrasting with the reduced uptake in MFN2-KO cells (**Fig. 4B**). Together, these findings indicate that MAM formation promotes the exogenous uptake and mobilization of cholesterol from the PM towards the ER, whereas MAM deficiency impairs this cholesterol flux.

**Fig. 4.**
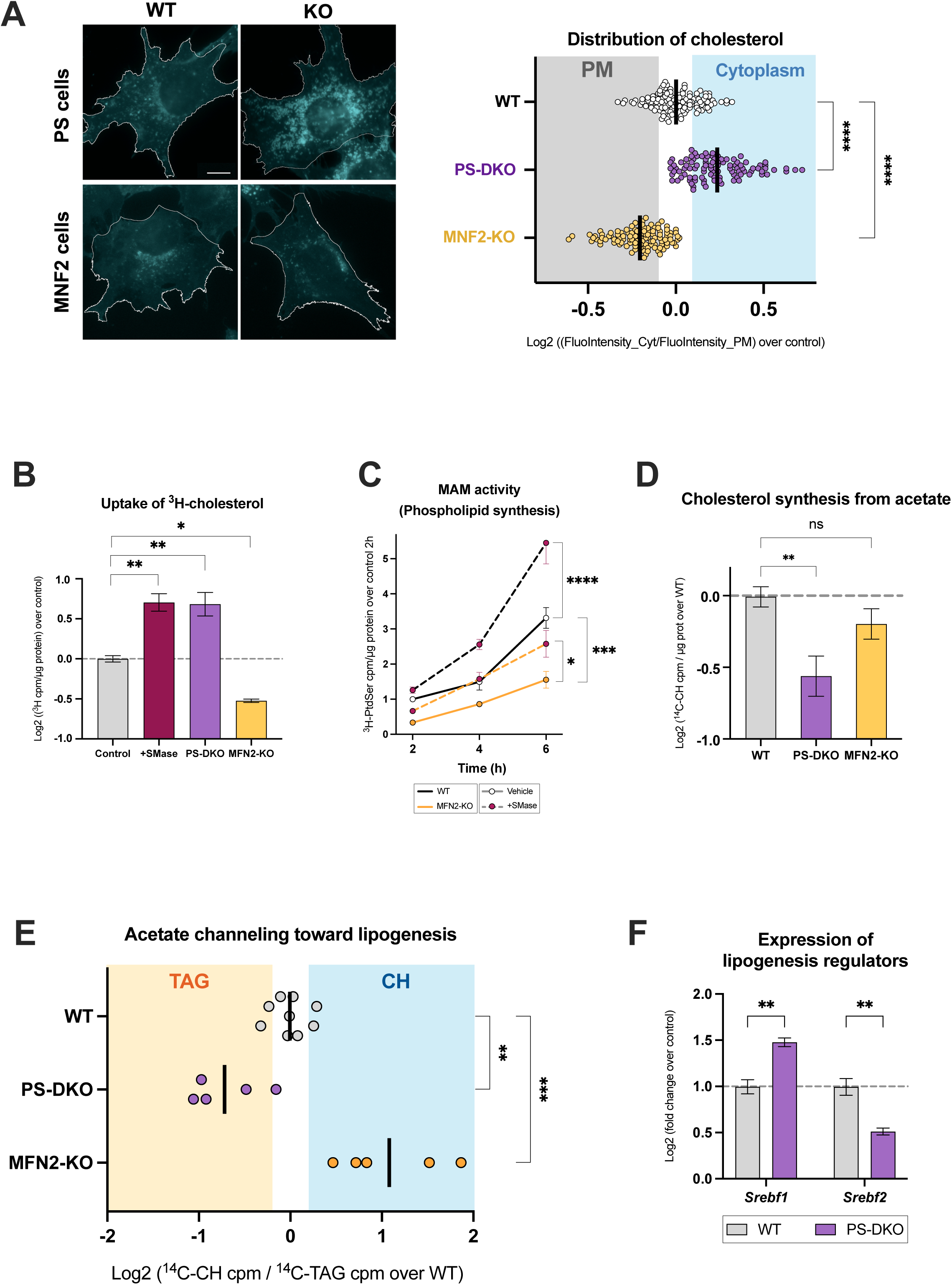
MAM formation induces cholesterol internalization and its delivery to the ER. **(A)** Representative images of PS-DKO or MFN2-KO MEFs stained with filipin and quantification of fluorescent signal compared to their controls. Scale bar = 10 μm (*n = 69-121 cells per condition*). **(B)** Quantification of exogenous ^3^H-cholesterol incorporation after 4h in 5% FBS media in the indicated cells compared to controls (*n = 3 biological replicates)*. **(C)** Measurement of MAM activity in MFN2-KO cells exposed to 1 U/mL SMase for the indicated times, assessed by phosphatidylserine (PtdSer) synthesis (*n = 3 independent experiments*). **(D)** Measurement of cholesterol biosynthesis from ^14^C-acetate in PS-DKO or MFN2-KO cells cultured in 10% FBS for 24h (*n = 5 biological replicates*). **(E)** Analysis of the incorporation of ^14^C-acetate into *de novo* synthesized cholesterol (CH) or triglycerides (TAG) in the indicated cells (*n = 5 biological replicates*). **(F)**. Quantification of mRNA levels of *Srebf1* and *Srebf2* (*n = 3 biological replicates*). All data are expressed as the mean ± SEM, * p<0.05, ** p<0.01, *** p<0.001, **** p<0.0001.

To stimulate cholesterol influx and its delivery to the ER, we exposed MAM-deficient cells to increasing concentrations of fetal bovine serum (FBS, containing cholesterol) **(Fig. S4B)** or exogenous cholesterol in the medium **(Fig. S4C)**. Compared to controls and PS-DKO cells, elevations in FBS concentration and cholesterol failed to boost cholesterol uptake and phospholipid synthesis in MFN2-KO cells. Conversely, treating these cells with SMase successfully mobilized cholesterol from the PM towards the ER, partially recovering MAM activity (**Fig. 4C**).

To further assess the contribution of MAM formation to cholesterol metabolism, we measured *de novo* cholesterol biosynthesis from ^14^C-acetate in these mutants, as described above. Consistent with our previous findings, PS-DKO cells, in which MAMs are constitutively upregulated, showed a significant reduction in cholesterol biosynthesis (**Fig. 4D**), accompanied by a shift in acetate incorporation into TAGs (**Fig. 4E**). This metabolic shift was further supported by reduced *Hmgcr* expression (**Fig. S4D**), reduced HMGCR activity (**Fig. S4E**) and decreased mRNA levels of *Srebf2*, together with increased *Srebf1* expression (**Fig. 4F**). Conversely, MAM-deficient MFN2-KO cells showed a significant increase in the expression and activity rate of HMGCR compared to controls (**Figs. S4D and S4E**), with rates of cholesterol synthesis from acetate similar to controls (**Fig. 4D**).

Together, these findings demonstrate a bidirectional crosstalk between the PM and MAM in the regulation of cholesterol homeostasis, whereby MAM formation regulates cholesterol and glycerolipid metabolism, redirecting acetate utilization toward TAG synthesis.

### The formation of MAM is induced by SR-B1-mediated uptake of cholesterol

Our data show that MAM formation represses the expression and processing of SREBP2, the master regulator of the expression of cholesterol biosynthesis enzymes and LDL-lipoprotein receptors, such as LDLR [27]. Therefore, it is possible that the stimulation of cholesterol uptake in cells where MAM is upregulated is a consequence of the sustained inhibition of SREBP2 pathways [20]. To determine if this is the case, we overexpressed a gene encoding the transcriptionally-active form of SREBP2 in PS-DKO cells, in which MAM is constitutively activated. However, we did not observe any effect on the internalization of cholesterol (**Figs. S5A and S5B**) or on the activity of the MAM-resident enzyme ACAT1 (**Fig. S5A**). This led us to conclude that MAM stimulates cholesterol internalization through a less-canonical, SREBP2-independent, pathway.

To explore this further, we quantified the gene expression of lipoprotein receptors sensitive to MAM modulation, by comparing cells exposed to SMase, PS-DKO mutants and MFN2-KO mutants to control cells. Our results show that the expression of genes encoding canonical lipoprotein receptors, such as *Ldlr* (low-density lipoprotein receptor) or *Lrp1* (LDL-related protein 1), were either reduced or not significantly changed by either the stimulation or inhibition of MAM formation (**Fig. 5A**). Conversely, the expression of the scavenger receptor B1 (SR-B1), known for its high affinity for HDL particles [17], was significantly increased in cells upon SMase treatment and in PS-DKO cells. Incubation of cells with SMase for 3h also stimulated the exposure of SR-B1 on the PM, without significantly affecting overall levels of this receptor, according to fluorescence microscopy data (**Fig. 5B**). On the other hand, PS-DKO cells displayed a reduction in SR-B1 at the PM despite presenting an elevation in the levels of this protein, as shown both by fluorescence microscopy (**Fig. 5B**) and surface biotinylation **(Fig. S5C)**. In MFN2-KO cells, however, the overall levels of SR-B1 were reduced without affecting its localization in the PM (**Fig. 5B**).

**Fig. 5.**
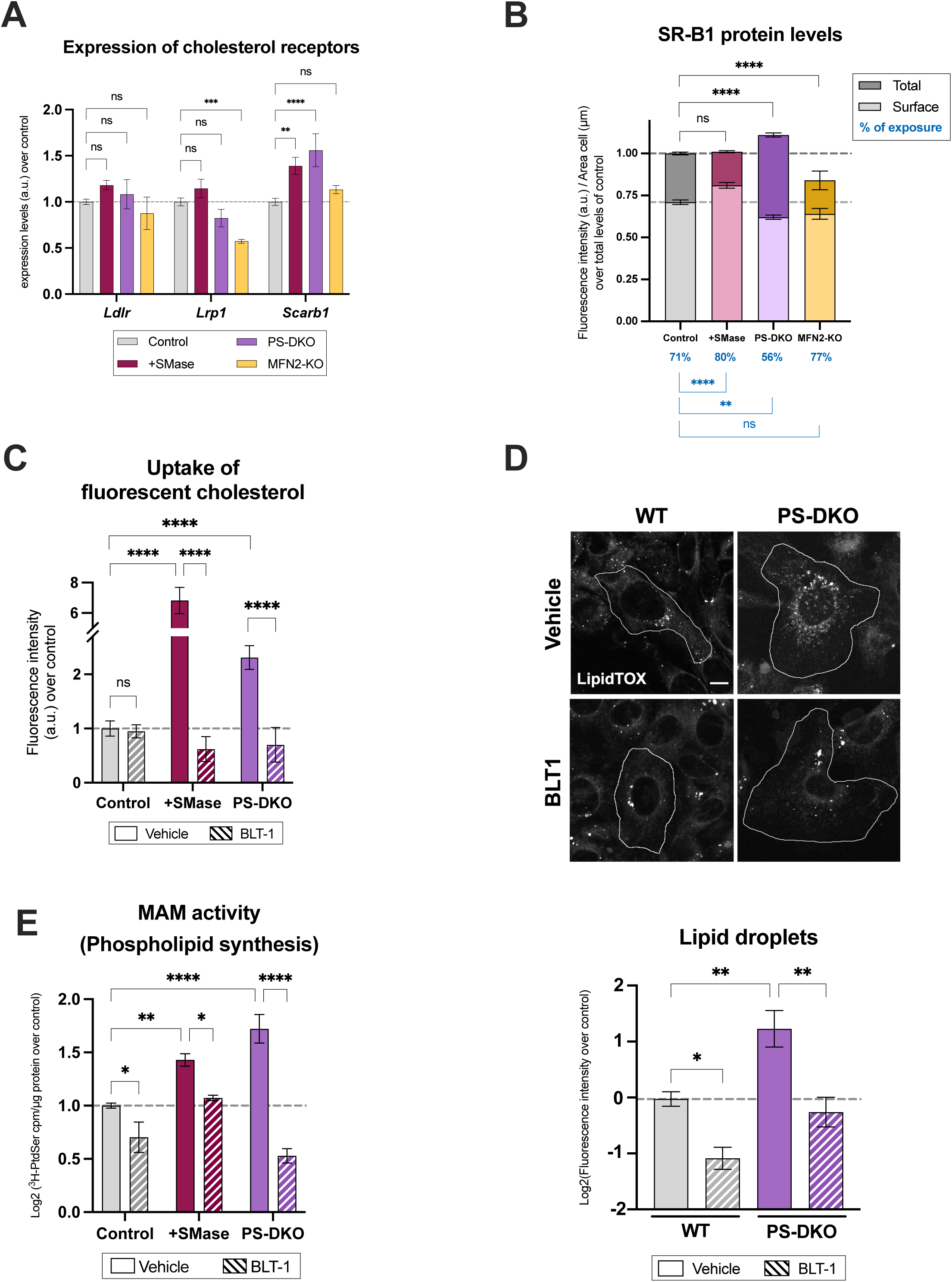
SR-B1-mediated uptake of cholesterol induces MAM formation. **(A)** Measurement of mRNA expression levels of the indicated lipoprotein receptors under the indicated conditions (*n = 3 biological replicates*). **(B)** Quantification of SR-B1 protein levels by immunofluorescence intensity in the indicated cells and conditions. The amount of protein exposed at the PM (Surface) over total levels (Total) was quantified, and was represented as the proportion of exposed SR-B1 (% of exposure) (*n = 51-250 cells per condition*). **(C)** Quantification of NBD-cholesterol incorporation into the indicated cells and conditions before and after incubation with 10 μM BLT-1 (*n = 3 biological replicates*). **(D)** Representative images of lipid droplet staining by LipidTOX in PS-DKO cells exposed to 10 μM BLT-1 or DMSO for 16h, scale bar = 10 μm. Quantification below (*n = 3 biological replicates*). **(E)** Measurement of MAM activity assessed by phosphatidylserine (PtdSer) synthesis in the indicated cells and conditions, after 4h in 5% FBS media (*n = 3 biological replicates*). All data are expressed as the mean ± SEM, * p<0.05, ** p<0.01, *** p<0.001, **** p<0.0001.

In light of these data, we next assessed SR-B1’s contribution to PM-MAM communication by incubating SMase-treated and PS-DKO cells with 10 µM BLT-1, an irreversible inhibitor of SR-B1 [28]. In both cell models, SR-B1 inhibition prevented the upregulation of cholesterol uptake (**Fig. 5C**), its esterification (**Fig. S5D**), and subsequent lipid droplet formation (**Fig. 5D**), without significantly affecting cholesterol efflux (**Fig. S5E**). Additionally, this treatment also effectively blocked the expected increase in cholesterol transport to the ER after boosting its levels in the PM by incubation with methyl-β-cyclodextrin-cholesterol complexes (**Fig. S5F).** SR-B1 inhibition was also able to rescue the upregulation of MAM in either PS-DKO or SMase-treated WT cells (**Fig. 5E**), the perinuclear distribution of cholesterol in PS-DKO cells (**Fig. S5G)** and the increased degree of ER-mitochondria apposition in the SMase-treated cells (**Fig. S5H**). Interestingly, SMase treatment also induced an increase in *Scarb1* expression that was abrogated by BLT-1 treatment (**Fig. S5I**), suggesting a feedback regulatory loop between SR-B1-dependent cholesterol uptake and MAM formation.

Taken together, our results suggest that cholesterol uptake via SR-B1 contributes to its delivery to the ER and the formation of MAM, compared to other canonical lipoprotein receptors.

### HDL-SR-B1 signaling induces MAM formation

Our results suggest that cholesterol internalization mediated by SR-B1 could contribute to the activation of MAM. However, evidence suggested that adding pure cholesterol straight to the medium of cultured cells, instead of feeding them with lipoproteins, can exacerbate the effect of scavenger receptors, and thus, overestimate the activation of SR-B1 in the uptake of cholesterol [29].

Therefore, we decided to evaluate MAM regulation and its impact on cholesterol metabolism under more physiological conditions, using different types of lipoproteins. To do that, we incubated WT cells with purified HDL, LDL, or VLDL over time and analyzed cholesterol distribution in the cell, relative to controls exposed to complete medium (containing 10% FBS). Compared to LDL or VLDL, exposure to HDL particles induced a significant redistribution of cholesterol from PM to the perinuclear areas (**Figs. 6A and S6A**), similar to those found in SMase-treated (**Fig. 2C**) or PS-DKO (**Fig. 4A**) cells. Notably, this redistribution of cholesterol was ameliorated in the presence of BLT1, confirming the relevance of SR-B1 to HDL uptake, but not to that of LDL or VLDL (**Fig. 6A**).

**Fig 6.**
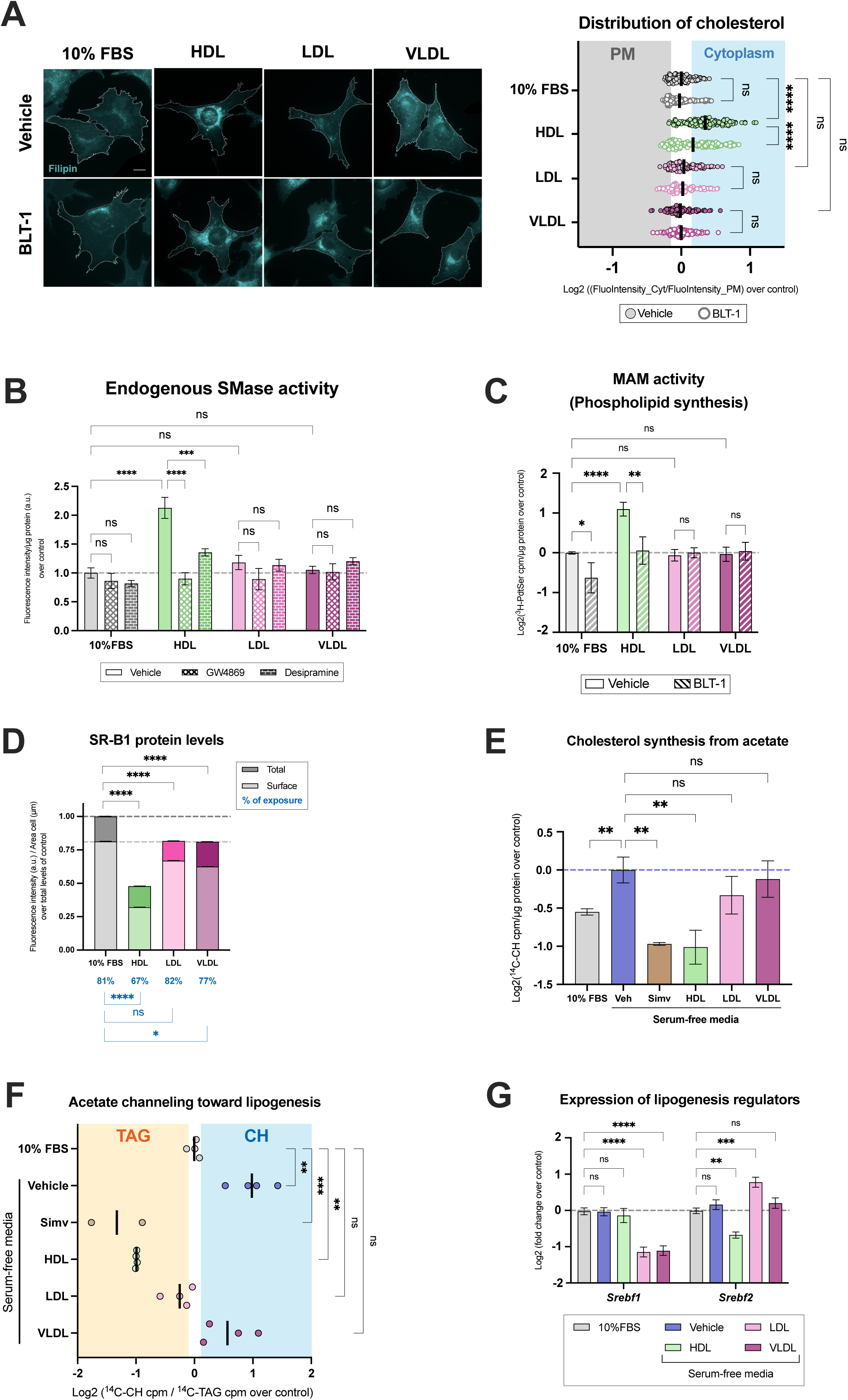
HDL-SR-B1 signaling induces MAM formation. All experiments were performed in MEF cells exposed to 5 ng/mL of different lipoproteins (HDL, LDL, VLDL) in serum-free media and indicated inhibitors for 6h. **(A)** Representative images of filipin staining of the distribution of cholesterol in cells exposed to the indicated lipoproteins before and after incubation with 10 μM BLT-1, scale bar = 10 μm Quantification at right (*n = 107-151 cells per condition*). **(B)** Measurement of endogenous SMase activity in cells exposed to 5 μM GW4869 or 10 μM desipramine (*n = 9 biological replicates*). **(C)** Measurement of MAM activity in cells incubated with the indicated lipoproteins before and after incubation with 10 μM BLT-1, assessed by phosphatidylserine (PtdSer) synthesis (*n = 14 biological replicates*). **(D)** Quantification of SR-B1 protein levels per cell, based on immunofluorescence intensity. The amount of protein exposed at the PM (Surface) over total levels (Total) were quantified, as well as the proportion of SR-B1 exposed (% of exposure) (*n = 68-134 cells per condition*). **(E)** Measurement of cholesterol synthesis *de novo* from ^14^C-acetate in serum-free cells exposed to 10% FBS, simvastatin (Simv) or lipoproteins (*n = 4 biological replicates*). (**F**) Analysis of the incorporation of ^14^C-acetate into *de novo* synthesized cholesterol (CH) or triglycerides (TAG) in the indicated cells, exposed to simvastatin (Simv) or the indicated lipoproteins for 6h (*n = 4 biological replicates*). **(G)** Measurement of mRNA expression levels of *Srebf1* and *Srebf2* (*n = 9 biological replicates*). All data are expressed as the mean ± SEM, * p<0.05, ** p<0.01, *** p<0.001, **** p<0.0001.

In agreement with this increase in cholesterol trafficking, exposure to HDL induced a significant activation of endogenous SMase activity over time (**Figs. 6B and S6B)** and the upregulation of MAM activities (**Fig. 6C**), compared to LDL or VLDL under the same conditions. As before, the impact of HDL on MAM activation was prevented by inhibition of SR-B1 by BLT-1 (**Fig. 6C**). Our data therefore indicate that MAM formation is preferentially stimulated by the SR-B1-mediated uptake of HDL particles, compared to the uptake of other lipoproteins of lower density.

Interestingly, these lipoproteins also had an impact on the levels and subcellular localization of SR-B1 as analyzed by fluorescence microscopy (**Fig. 6D**). Specifically, HDL-treated cells displayed overall decreased SR-B1 levels, whereas LDL and VLDL did not affect the exposure of SR-B1 in the PM (**Fig. 6D**) but stimulated the expression of *Ldlr* (**Fig. S6D**).

HDL incubation also induced a significant downregulation of the incorporation of ^14^C-acetate into cholesterol, similar to cells treated with the HMGCR inhibitor simvastatin, whereas incubation with LDL and VLDL particles had the opposite effect (**Figs. 6E and S6E**). In agreement, we also observed a significant reduction in the expression of *Srebf2* in HDL-treated cells, although there were no significant changes in mRNA levels of *Hmgcr* and most pre-squalene enzymes (**Fig. S6F**). As in the case of SMase-treated and PS-DKO cells, this significant reduction in cholesterol synthesis in cells incubated with HDL was associated with the shunting of labeled acetate into TAGs (**Fig. 6F**), despite not presenting significant changes in *Srebf1* expression (**Fig. 6G**).

On the other hand, exposure to LDL or VLDL showed no significant effect on the reduction of cholesterol biosynthesis, compared to cells in serum-free media (**Fig. 6E**). However, while serum-free cells displayed remarkable acetate shunting towards cholesterol synthesis, cells exposed to LDL or VLDL seem to keep the homeostatic balance of cells in media containing 10% FBS (**Fig. 6F**). Interestingly, this LDL and VLDL phenotype was accompanied by reduced *Srebf1* expression and increased *Srebf2* expression (**Fig. 6G**).

These results suggest that, compared to LDL and VLDL lipoproteins, MAM formation is particularly stimulated by the uptake of HDL particles, inducing a faster inhibition of cholesterol biosynthesis and the channeling of acetyl-CoA substrates towards glycerolipid synthesis.

Altogether, our data indicate that the formation of MAM domains acts as a cholesterol-sensing mechanism in the ER, reflecting changes in cholesterol levels in the PM and orchestrating feedback reactions to help maintain cholesterol homeostasis (**Fig. 7**). We also show that MAM is sensitive to the extracellular lipoprotein environment, thereby contributing to the regulation of cellular metabolic flexibility.

**Fig. 7.**
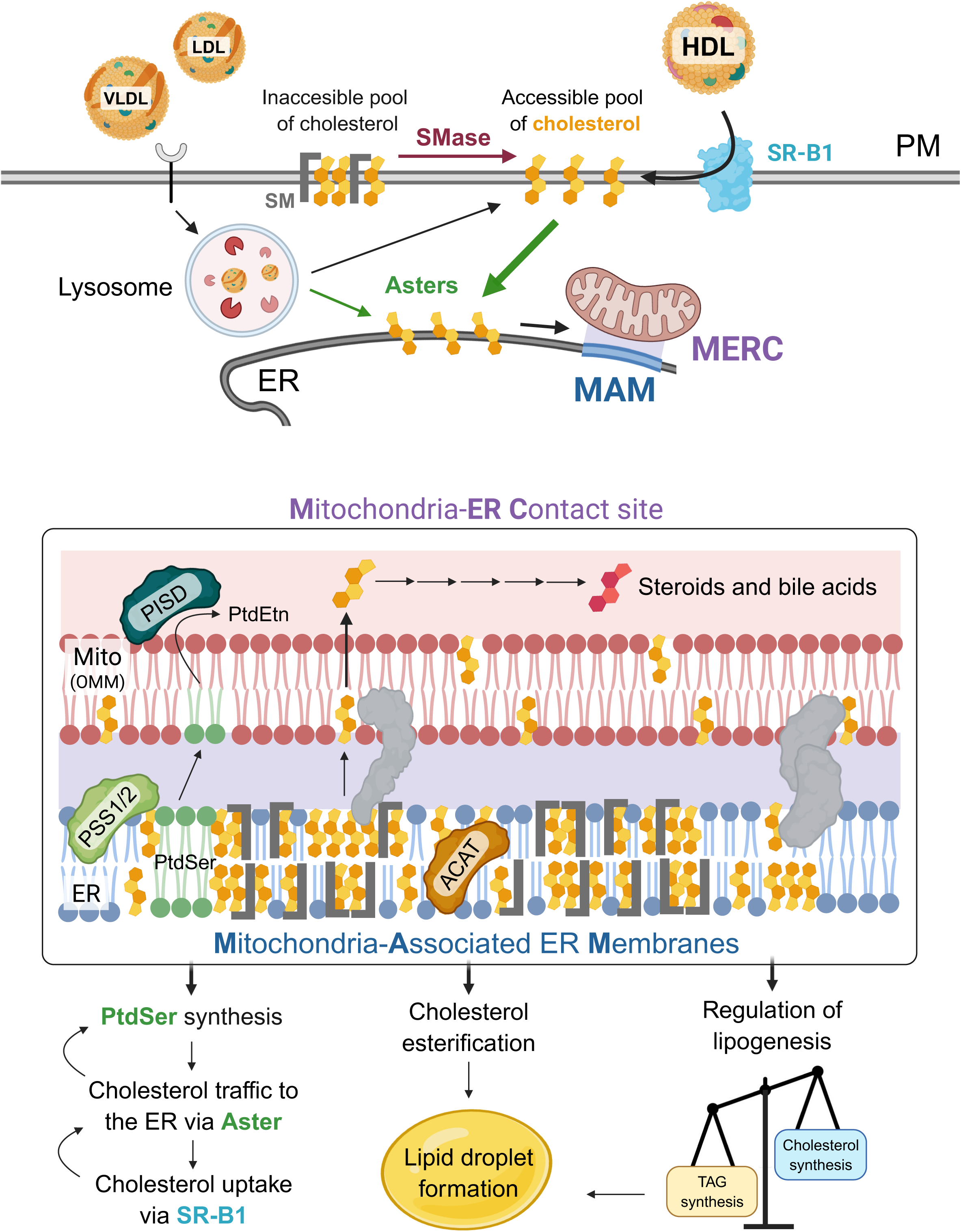
MAM formation integrates cellular responses to cholesterol availability. We propose a model in which the SRB1-HDL-MAM axis is a mechanism to coordinate metabolic responses to increased cholesterol availability in the cell. Increases in the accessible pool of cholesterol at the plasma membrane (PM) is transferred to the ER membrane, in part through the Aster protein family. At the ER, cholesterol can promote the formation of cholesterol-rich membrane domains characteristic of MAM, facilitating the recruitment of specific lipids and proteins and promoting their association with the outer mitochondrial membrane (OMM) to establish mitochondria–ER contact sites (MERCs). Importantly, compared with other lipoproteins, HDL exposure strongly promotes cholesterol trafficking and MAM formation, compared to LDL and VLDL.

## DISCUSSION

In this work, we show that MAM domains and their association with mitochondria are dynamically induced by the mobilization of accessible cholesterol from the PM to the ER, rather than by cholesterol biosynthesis per se. In fact, MAM formation is accompanied by the suppression of cholesterol biosynthesis and by increased channeling of fatty acids toward cholesterol esterification, consistent with a coordinated response to restore cellular lipid homeostasis. Moreover, we demonstrate that this response is particularly stimulated by the SR-B1-HDL uptake, establishing a functional link between extracellular lipoprotein-derived cholesterol, intracellular cholesterol trafficking, and MAM formation (**Fig. 7**). These findings suggest that MAMs should be viewed not simply as structural ER–mitochondria contact sites, but as dynamic lipid-regulated domains that participate in the cellular response to changes in cholesterol availability.

A *primary* implication of this study is the delineation of a distinct mechanism driving the formation of MAMs and MERCs.

MAMs have emerged as important sites for the coordination of multiple metabolic processes, including lipid synthesis, cholesterol metabolism, calcium signaling, and mitochondrial function [2, 30]. Much of the work defining ER–mitochondria contacts, however, has focused on the regulation of artificial or endogenous protein tethers.

Our data provide an alternative perspective by which changes in membrane lipid composition can themselves drive the formation of MAM domains and their association with mitochondria – by the formation of a cholesterol-enriched, raft-like domain. As such, MAM formation would transiently partition the ER membrane and selectively concentrate or exclude proteins according to their lipid environment, thereby creating a localized platform in which multiple metabolic enzymes can be coordinately regulated in response to extracellular cholesterol.

It is also possible that, within the ER, cholesterol may exist in distinct functional pools, each with different regulatory properties to influence metabolic pathways. Therefore, MAM pools may also coexist with other domains and organelle contact sites, contributing to the orchestration and spatial organization of cellular metabolism.

In agreement with this perspective, we show that the formation of MAM represses acetate use for cholesterol synthesis, channeling it into TAGs. Importantly, acetate is selectively converted into specific functional pools of acetyl-coenzyme A (acetyl-CoA) that participate in central carbon metabolism for membrane generation, energy production, or epigenetic signaling. Our data thus indicate that the formation of the MAM contributes to the compartmentalization of these lipogenesis pathways. Similarly, through contact sites with mitochondria, the MAM can also modulate the shunting of cholesterol for the synthesis of steroids in mitochondria [31, 32]. Moreover, MAM formation links cholesterol distribution and the modulation of mitochondria and cellular bioenergetics [33].

A central question that remains unresolved is how cholesterol-induced MAM domains engage mitochondria. Although our data establish a clear relationship between cholesterol mobilization, MAM formation, and increased ER–mitochondria association, we did not determine whether mitochondrial recruitment is driven primarily by changes in membrane physical properties or by the recruitment or activation of specific ER–mitochondria tethering proteins. Defining this mechanism will be important for understanding how lipid sensing at the ER is coupled to MAM biogenesis and mitochondrial function.

A *second* implication of our findings is that MAMs may represent a functionally regulatory pool of cholesterol in the ER. Cellular cholesterol homeostasis is controlled through tightly integrated pathways of synthesis, uptake, transport, efflux, and esterification, largely coordinated by master lipid regulators such as SREBP-1 and 2 [11].

Recent evidence has highlighted that, in addition to these complex pathways, total cellular or organellar cholesterol content is less informative than the size and chemical activity of specific cholesterol pools, and their accessibility to the cholesterol-sensing machinery.

For example, it is well established that the ER contains a relatively small amount of cholesterol, but it forms pools of active cholesterol essential for sensing the cell’s cholesterol status, and inducing feedback responses [10, 34]. Moreover, multiple reports demonstrate that these cholesterol pools are highly stimulated by treatment with SMase [35, 36, 37]. However, the identity and regulation of these pools of cholesterol in the ER are not completely understood.

Our findings raise the possibility that MAM represents one such specialized regulatory cholesterol pool within the ER that can sense cholesterol levels in the PM and contribute to the compartmentalization and downregulation of cholesterol biosynthesis. In agreement, previous reports have shown that MAM is enriched in enzymes related to the post-squalene pathway of cholesterol biosynthesis [38].

Our data also suggest that the pool of PM cholesterol that induces MAM domains in the ER is regulated by Asters. Interestingly, previous data indicate that the delivery of cholesterol through this non-vesicular mechanism requires phosphatidylserine, a phospholipid synthesized solely at the MAM *space* [37].

MAM formation could therefore establish a local cholesterol-sensing environment in which increased cholesterol availability suppresses further synthesis while promoting its esterification or transfer into alternative metabolic pathways.

A *third* important finding is that MAM formation is particularly responsive to HDL-mediated cholesterol uptake. Although LDL represents the classical pathway for cellular cholesterol delivery, increasing evidence indicates that HDL-SR-B1 pathways represent an essential route for cholesterol flux and metabolic regulation [39]. In our experiments, HDL exposure produced greater cholesterol internalization and stronger MAM activation than did LDL or VLDL, and was associated with a more pronounced suppression of *de novo* cholesterol biosynthesis.

HDL and LDL may differentially influence MAM formation because they deliver cholesterol to the cell through distinct pathways and with different kinetics. It is well known that HDL-SR-B1-mediated cholesterol uptake promotes rapid, concentration-dependent cholesterol transfer, which may facilitate more rapid mobilization of accessible cholesterol from the PM to the ER than LDL-derived cholesterol can [17, 40]. Consistent with this possibility, previous studies have shown that, whereas LDL-derived cholesterol is initially delivered to the PM before reaching the ER, HDL-derived cholesterol could be delivered directly to the ER [41]. Thus, the higher rate of cholesterol transfer associated with SR-B1 may provide a stronger or more rapid signal for MAM formation and the subsequent suppression of cholesterol biosynthesis. Importantly, this internalization pathway does not exclude the possibility that LDL can ultimately induce similar responses, but rather suggests that the kinetics and intracellular routing of cholesterol delivery may determine the extent and timing of MAM activation.

The different effects of HDL- and LDL-derived cholesterol may also reflect the distinct PM cholesterol pools that are mobilized due to the heterogeneity of lipoprotein particles themselves. HDL particles vary substantially in size, lipid composition, and associated proteins, and their effects on cellular metabolism may therefore depend on particle subtype rather than on lipid content alone. Hence, rapid HDL-SR-B1-mediated cholesterol transfer could replenish and promote the mobilization of specific accessible PM cholesterol pools. In contrast, LDL may reach the ER through slower or distinct trafficking routes and could generate different cholesterol-containing domains with distinct effects on the cholesterol biosynthetic machinery and on SREBP2 signaling. Thus, the formation of MAMs may depend not simply on the amount of cholesterol delivered to the cell, but on the rate, route, and membrane pool through which cholesterol is mobilized. Moreover, other receptors and trafficking pathways may also contribute to this response, and the potential involvement of additional ER-associated contact sites, including PM- and Golgi-associated ER membranes, should not be excluded [42, 43].

Together, our findings support a model in which MAM is significantly induced by the HDL–SR-B1 cholesterol pathway (**Fig. 7**), providing a mechanism for adapting cellular metabolism to different environmental conditions.

## MATERIALS AND METHODS

### Reagents

All reagents, equipment and catalog numbers are listed in the Resources Table.

### Cells

All assays were performed in WT mouse embryonic fibroblasts (MEFs). *Psen1/2*-double knock-out (called PS-DKO) MEFs were obtained from Dr. Bart De Strooper (University of Leuven). *Mfn2*-knock-out (MFN2-KO) MEFs, as well as the relevant control MEFs, were kind gifts from Dr. David Chan (California Institute of Technology). In some specified experiments, results were validated in SH-SY5Y cells, obtained from the American Type Culture Collection.

### Cell treatments

Cells were incubated with the indicated cholesterol sources in serum-free high-glucose DMEM without prior lipid starvation, unless otherwise specified. Cells were treated with 2.5 μCi/mL [^3^H]-cholesterol complexed with 2% BSA, 2.5 μCi/mL [^3^H]-cholesterol complexed with methyl-β-cyclodextrin, 1 μM NBD-CH, or 5 ng/mL purified lipoproteins (HDL, LDL, or VLDL) for the indicated times.

Plasma membrane-to-ER cholesterol trafficking was stimulated by treatment with 1 U/mL SMase for 3 h in media containing 5% FBS. Unless otherwise indicated, pharmacological inhibitors were added 15 min before treatment, with DMSO used as the vehicle control. SR-B1 activity was inhibited using 10 μM BLT1. Neutral and acidic SMase activities were inhibited with 5 μM GW4869 and 10 μM desipramine, respectively. HMGCR was inhibited with 20 μM atorvastatin or 50 nM simvastatin. Intracellular cholesterol trafficking was inhibited with 1 μM U18666A or 2 μM AI-d3.

### Subcellular fractionation

Purification of ER, crude membranes (CM), mitochondria and MAM was performed and analyzed as previously described [44].

### Western blotting

After quantification of sample protein concentrations via the BCA assay, equal amounts of protein were boiled in Laemmli sample buffer and run in 4-20% Tris-glycine gels. Proteins were detected using the antibodies listed in the Reagents Table.

### Quantitative reverse transcription–polymerase chain reaction (qRT–PCR)

Total RNA was extracted using TRIzol® Reagent according to the manufacturer’s instructions and was quantified by NanoDrop2000. One microgram of total RNA was used to generate cDNA by RT–PCR using the High-Capacity cDNA Reverse Transcription Kit. Quantitative Real-Time PCR was performed in triplicate in a StepOnePlus™ Real-Time PCR System. The expression of each gene under study was analyzed using specific predesigned TaqMan Probes (see Reagents Table) and normalized against 18S rRNA expression as an internal control.

### Phospholipid transfer

2.5 µCi/mL ^3^H-serine was added to the cells for the specified time-points. After extensive washing, cells were collected and lipids extracted and separated by TLC as detailed in [23].

### Analysis of ER-mitochondria apposition

Cells were co-transfected with mCherry-SEC61B (Addgene plasmid, #90994) and mitoGFP (Addgene plasmid, #159096) at a 1:1 ratio using Lipofectamine™ 2000 Transfection Reagent in serum-free DMEM for 4-6h following manufacturer instructions. 16h post-transfection, cells were treated with SMase and/or BLT-1 for 1h or 3h. Cells were fixed with 4% paraformaldehyde and, after extensive washing, coverslips were mounted with Fluoromount-G^TM^. Images of double-transfected cells were acquired by confocal microscopy, and ER-mitochondria apposition was analyzed as in [45].

### 3H-cholesterol incorporation and esterification assays

Cholesterol incorporation and esterification were assessed as in [26]. Cells were incubated with 2.5 µCi/mL of ^3^H-cholesterol in serum-free DMEM supplemented with 2% fatty acid-free BSA and allowed to equilibrate at 37°C for at least 30 minutes. In selected experiments, the same amount of ^3^H-cholesterol was complexed with methyl-β-cyclodextrin and then administered to the cells for the specified durations. Afterward, cells were rinsed and harvested in PBS. For some experiments, subcellular fractionation was performed prior to lipid extraction. Equal protein amounts were used to extract lipids by using three volumes of chloroform:methanol (2:1 v/v). Following vortexing and centrifugation at 8,000 g for 5 minutes, the organic phase was dried under nitrogen. The resulting lipid residue was reconstituted in 20 µL of chloroform and applied to a thin-layer chromatography (TLC) plate alongside unlabeled standards. A solvent mixture of hexanes/diethyl ether/acetic acid (80:20:1 v/v/v) was used for TLC. Iodine-stained bands corresponding to cholesterol and cholesteryl esters were scraped off and measured in a scintillation counter.

### Cholesterol and lipid droplets staining

Cholesterol uptake was assessed by fluorescence microscopy following incubation with 1 μM NBD-CH as in [26]. For filipin staining, cells were fixed with 4% paraformaldehyde and incubated with 50 μg/mL filipin for 2h at room temperature. After extensive washing, coverslips were mounted with ProLong™ Diamond Antifade and imaged by epifluorescence microscopy.

Lipid droplets were stained with HCS LipidTOX™ Green according to the manufacturer’s instructions. Following extensive washing, coverslips were mounted with Fluoromount-G™ and imaged by confocal fluorescence microscopy.

Fluorescence intensity was quantified using ImageJ. Raw fluorescence intensity within manually defined regions of interest (ROIs) was normalized to the corresponding cell area. ROIs were defined either around individual cells or specific cellular regions (PM, cytoplasm), as indicated.

### Cholesterol efflux

Cells were loaded with 2.5 μCi/mL of ^3^H-cholesterol prepared as described above. Three hours after incubation, cells were washed to remove excess exogenous radioactive cholesterol and incubated in unlabeled media in the presence or absence of the indicated inhibitors. After the indicated post-incubation times, media was recovered, briefly centrifuged at 1,500 g for 5 min to remove any debris, transferred to scintillation vials containing 5 mL of scintillation cocktail and radioactivity was measured in a scintillation counter.

### HMG-CoA reductase (HMGCR) activity assay

HMGCR activity was measured following manufacturer instructions (Sigma; #CS1090) from equal protein amounts.

### Cholesterol and TAGs synthesis *de novo* from acetate

Cells were incubated with 0.5 µCi/ mL ^14^C-acetate and the co-treatments at the times specified in each experiment. Cells were rinsed and harvested in PBS with protease inhibitors. A small aliquot was saved for protein quantification. Lipids were extracted by using three volumes of chloroform:methanol (2:1 v/v). Following vortexing and centrifugation at 8,000 g for 5 minutes, the organic phase was dried under nitrogen. The resulting lipid residue was reconstituted in 20 µL of chloroform and applied to a thin-layer chromatography (TLC) plate alongside unlabeled standards. A solvent mixture of hexanes/diethyl ether/acetic acid (80:40:1.5 v/v/v) was used for TLC. Radioactive signal of iodine-stained bands corresponding to cholesterol and TAG were measured by a TLC reader or manually scraped and measured in a liquid scintillation counter. Radioactive signal data were normalized by the amount of protein per sample.

### Sphingomyelinase activity assay

Cells were incubated with 1 μM C16-sphingomyelin-Cy5 at 37°C for the indicated times. Following extensive washing, cells were collected and extracted with chloroform:methanol:H₂O (2:1:1). The organic and aqueous phases were separated by centrifugation at 8,000 × g for 5 min. The aqueous phase, containing fluorescent phosphocholine generated by sphingomyelin hydrolysis, was collected, and fluorescence was measured using a plate reader (excitation/emission, 645/685 nm).

### Transient expression of the mature form of SREBP2

PS-DKO cells were transfected with the truncated mature form of SREBP2 (2xFLAG-SREBP2, aa 1-482, Addgene plasmid #26807), which is targeted to the nucleus where it activates cholesterol biosynthesis gene programs. 16h post-transfection, cells were assayed for cholesterol uptake and esterification, as well as protein expression of 2xFLAG-SREBP2.

### Labeling and purification of plasma membrane proteins

Plasma membrane proteins were biotinylated by incubation with 1 mg/mL of EZ-Link™ Sulfo-NHS-LC-Biotin in labeling buffer (20 mM HEPES, 150 mM NaCl, 2 mM CaCl_2_, pH 8.5) at 4°C for 45 minutes under dark conditions. Subsequent washing steps with PBS containing glycine were conducted to quench unreacted biotin. Cells were then lysed in 50 mM Tris pH 7.5, 5 mM EGTA, 10 mM EDTA supplemented with protease inhibitors by sonication and addition of 1% sodium deoxycholate. After centrifugation, the supernatant containing the lysate was collected, and a fraction was reserved as input. Streptavidin-coated beads, pre-washed in 50 mM Tris pH 7.5, were added to the lysate and incubated overnight at 4°C. Following extensive washing, the bound proteins were eluted by heating at 45°C in 1xLaemmli buffer to prevent protein aggregation.

### SR-B1 exposure by fluorescence microscopy

After the indicated treatments, cells were fixed with 4% paraformaldehyde for 15 min. Cells corresponding to “Total SR-B1” were permeabilized with 0.1% Triton X-100 for 10 min, while “surface SR-B1” cells were not permeabilized. Fixed cells were blocked with 5% BSA for 1h, following an incubation with SR-B1 antibody in 5% BSA overnight at 4°C and a secondary incubation with anti-rabbit Alexa 647 antibody. After extensive washes, coverslips were mounted with ProLong^TM^ antifade and visualized by epifluorescence microscopy.

Images were quantified using ImageJ. Reported fluorescence intensity represents the raw intensity of the region of interest (ROI) normalized by cell area. Analysis was restricted to manually defined ROIs, representing individual cells.

## Statistical analysis

Data represent mean ± SEM. Normal distribution of data was determined using the Shapiro-Wilk’s test when possible. The statistical analysis was performed using GraphPad Prism v10.3.1. The Brown-Forsythe test was used to compare variance between groups. Statistical significance was determined by a two-tailed t-test, 1-way ANOVA or 2-way ANOVA, followed by the Tukey’s or Sidak’s multiple comparison post-hoc test. For filipin distribution and acetate channeling graphs, the control window corresponds to the control group mean ± 1SD. Values of p < 0.05 were considered statistically significant. * p<0.05, ** p<0.01, *** p<0.001, **** p<0.0001. The investigators were not blinded when quantifying or collecting data of imaging experiments.

## Supporting information

Reagents and antibodies

## Acknowledgements

This work was supported by the Spanish Ministry of Science, Innovation and Universities (PID2021-126818NB-I00 to EA-G; FPU fellowship FPU22/00245 to MU; FPI fellowship PRE2022-104771 to ACF; CAM fellowship PIPF-2024/SAL-GL-34942 to LSC, co-financed by The European Social Fund Plus; FPU fellowship FPU23/02047 to CA-B), Comunidad de Madrid (2024-T1SAL-GL-31373 Talento César Nombela to JM), the U.S. National Institutes of Health (R01-AG056387-01 to EA-G; T32-DK007647 to RRA and KAT; R01-NS117538 to EAS) and the U.S. Department of Defense (FA9550-11-C-0028 to RRA).

## Author Contributions Statement

Conceived the project: EA-G, EAS and JM. Designed experiments: EA-G, EAS, JM and MU. Generated experimental data: MU, EA-G, JM, KK, ACF, LSC, MP, NGL, DL, RRA, CA-B, KAT, TDY and KRV. Wrote the manuscript: EA-G and MU. Critically edited the manuscript: all authors. Approved final version of the manuscript: all authors.

---

**Fig S1.**
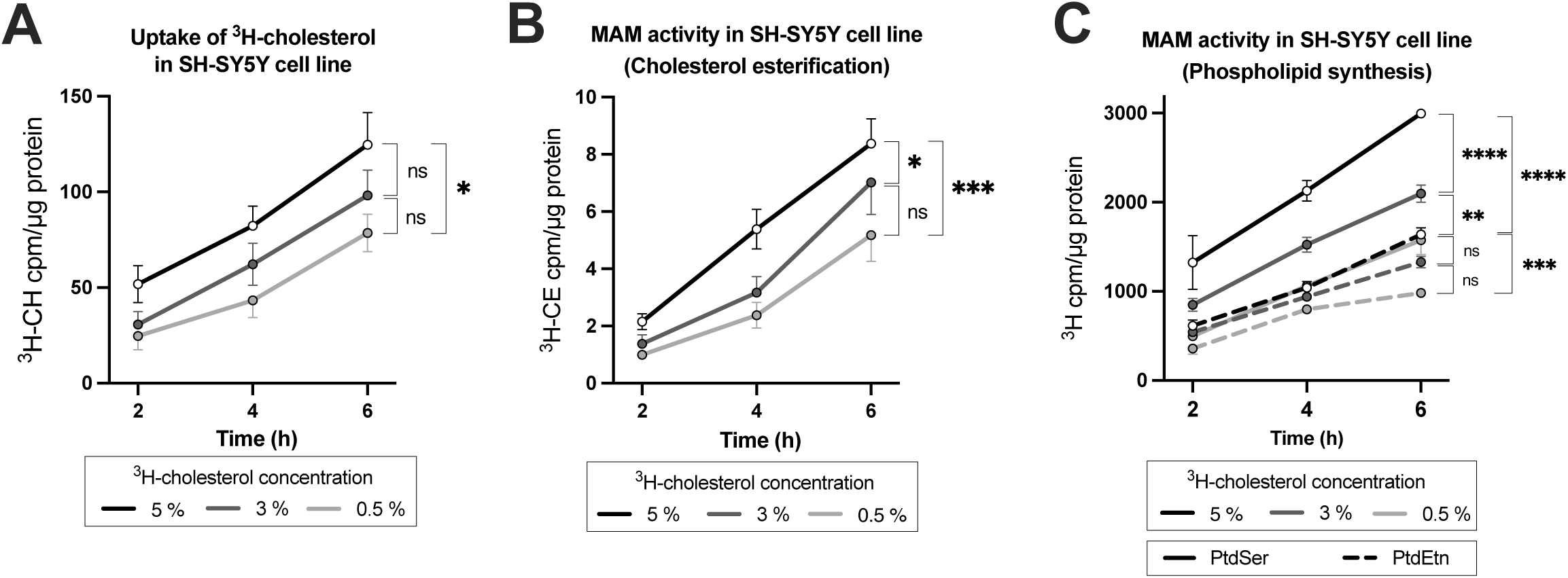
Cholesterol internalization induces MAM formation. **(A)** Quantification of exogenous ^3^H-cholesterol internalization into SH-SY5Y cells at increasing concentrations and times of incubation (*n = 3 biological replicates*). **(B-C)** Measurement of MAM activity in SH-SY5Y cells exposed to increasing concentrations of exogenous ^3^H-cholesterol, assessed by **(B)** cholesterol esterification (CE) and **(C)** phosphatidylserine (PtdSer) and phosphatidylethanolamine (PtdEtn) synthesis (*n = 3 biological replicates*). All data are expressed as the mean ± SEM, * p<0.05, ** p<0.01, *** p<0.001, **** p<0.0001.

**Fig S2.**
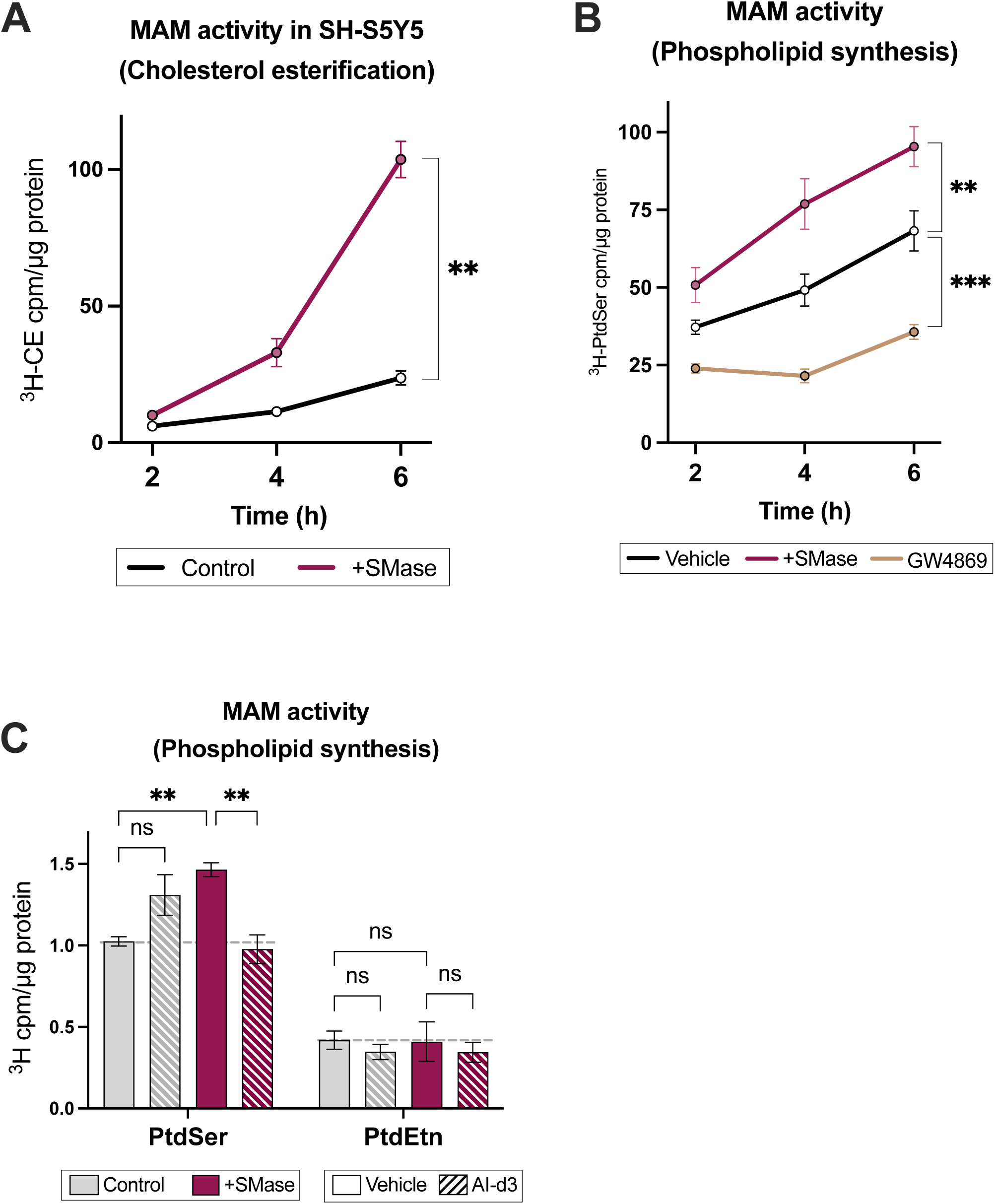
Cholesterol delivery to the ER induces MAM formation. **(A)** Measurement of MAM activity in SH-S5Y5 cells exposed to 1U/mL SMase in 5% FBS media over time, assessed by cholesterol esterification (CE) formation (*n = 3 biological replicates*). **(B)** Measurement of MAM activity in MEF cells exposed to 1U/mL SMase or 5 μM GW4869 over time in 5% FBS media, assessed by phosphatidylserine (PtdSer) synthesis (*n = 3 biological replicates*). **(C).** Measurement of MAM activity in MEF cells exposed to 1U/mL SMase and 2 μM AI-d3 for 3h in 5% FBS media, assessed by phosphatidylserine (PtdSer) and phosphatidylethanolamine (PtdEtn) synthesis (*n = 3 biological replicates*). All data are expressed as the mean ± SEM, ** p<0.01, *** p<0.001.

**Fig S3.**
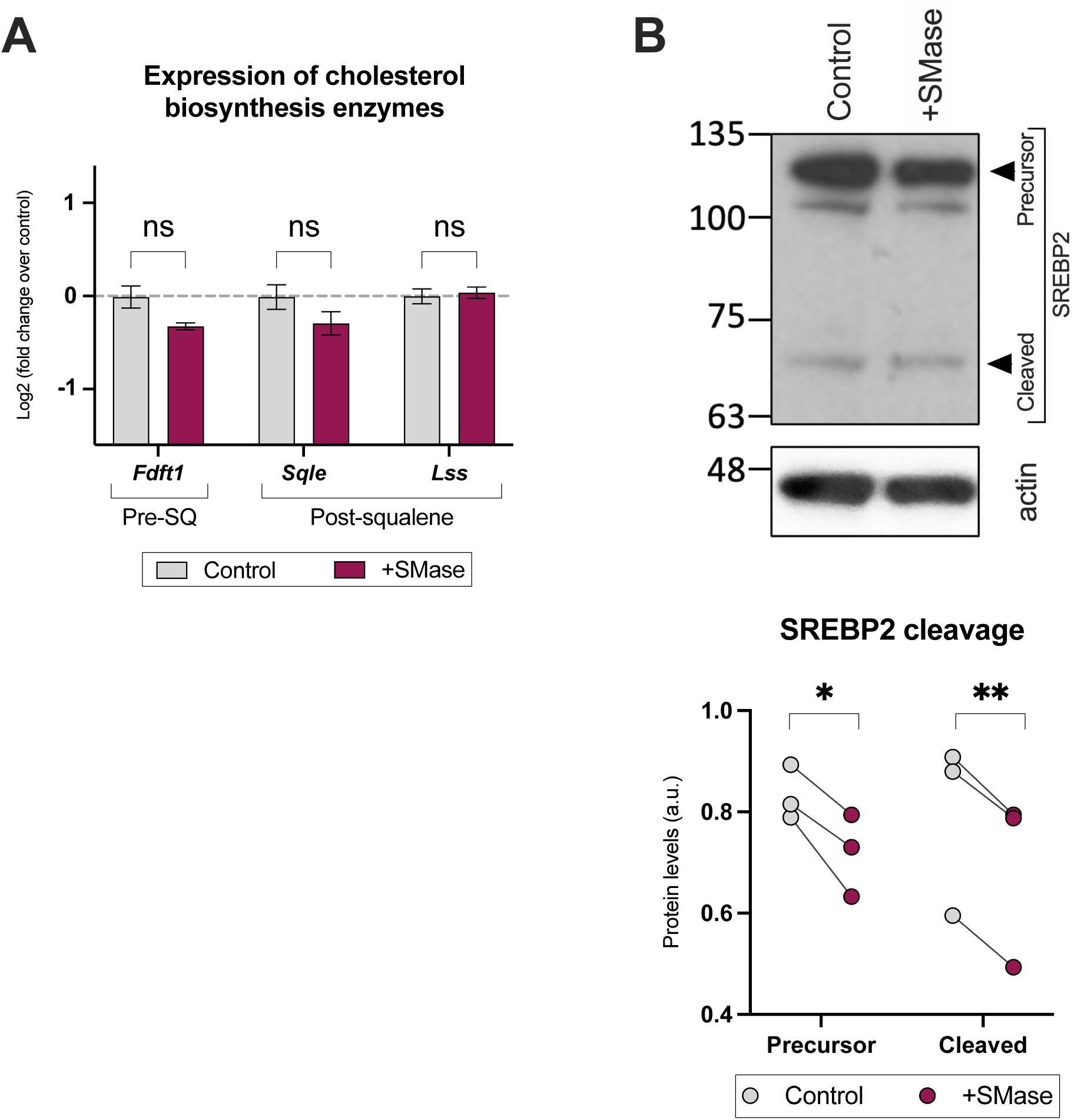
Cholesterol delivery to the ER represses cholesterol biosynthesis. All experiments were performed in MEF cells exposed to 1U/mL SMase for 3h in 5% FBS media. **(A)** Measurement of mRNA expression levels of cholesterol biosynthesis enzymes, pre- and post-squalene (SQ) (*n = 3 biological replicates*). **(B)** Representative image of western blot against SREBP2, showing the precursor and cleaved form. Sizes at left in kDa. Below, quantification of protein levels normalized to the loading control (β-actin) (*n = 3 biological replicates*). All data are expressed as the mean ± SEM, * p<0.05, ** p<0.01.

**Fig S4.**
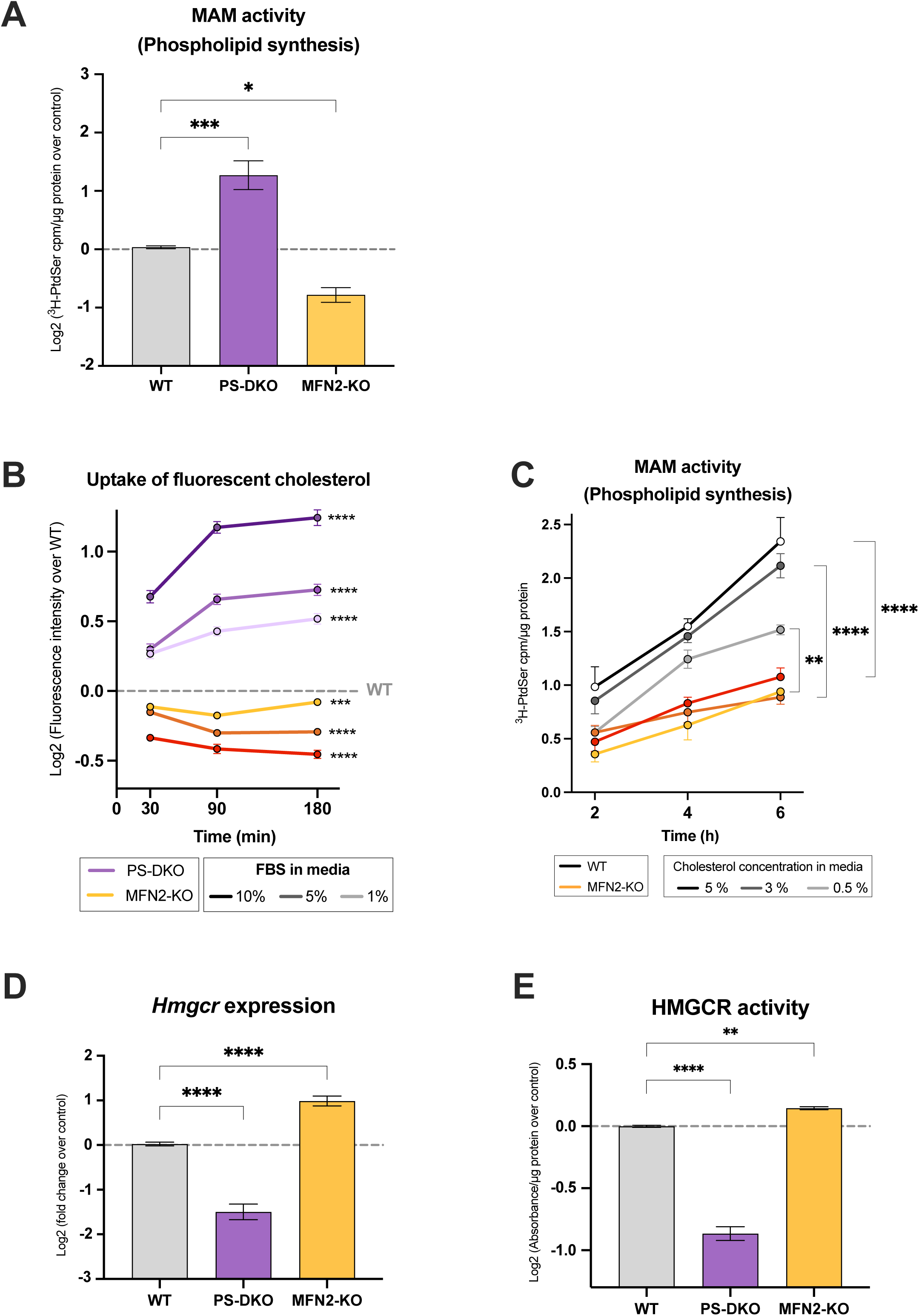
MAM formation induces cholesterol internalization and its delivery to the ER. **(A)** Measurement of MAM activity in PS-DKO or MFN2-KO cells, assessed by phosphatidylserine (PtdSer) synthesis over 5h (*n = 9 biological replicates*). **(B)** Quantification of NBD-cholesterol internalization by fluorescence intensity in PS-DKO or MFN2-KO over control WT at the indicated times (*n = 32 cells per condition)*. **(C)** Measurement of MAM activity in MFN2-KO cells exposed to the indicated concentrations of exogenous cholesterol, assessed by phosphatidylserine (PtdSer) synthesis over time (*n = 3 biological replicates*). **(D)** Measurement of mRNA expression levels of *Hmgcr* in PS-DKO or MFN2-KO cells (*n = 3 biological replicates*). (**E)** Measurement of HMGCR activity in PS-DKO or MFN2-KO cells (*n = 3 biological replicates*). All data are expressed as the mean ± SEM, * p<0.05, ** p<0.01, *** p<0.001, **** p<0.0001.

**Fig S5.**
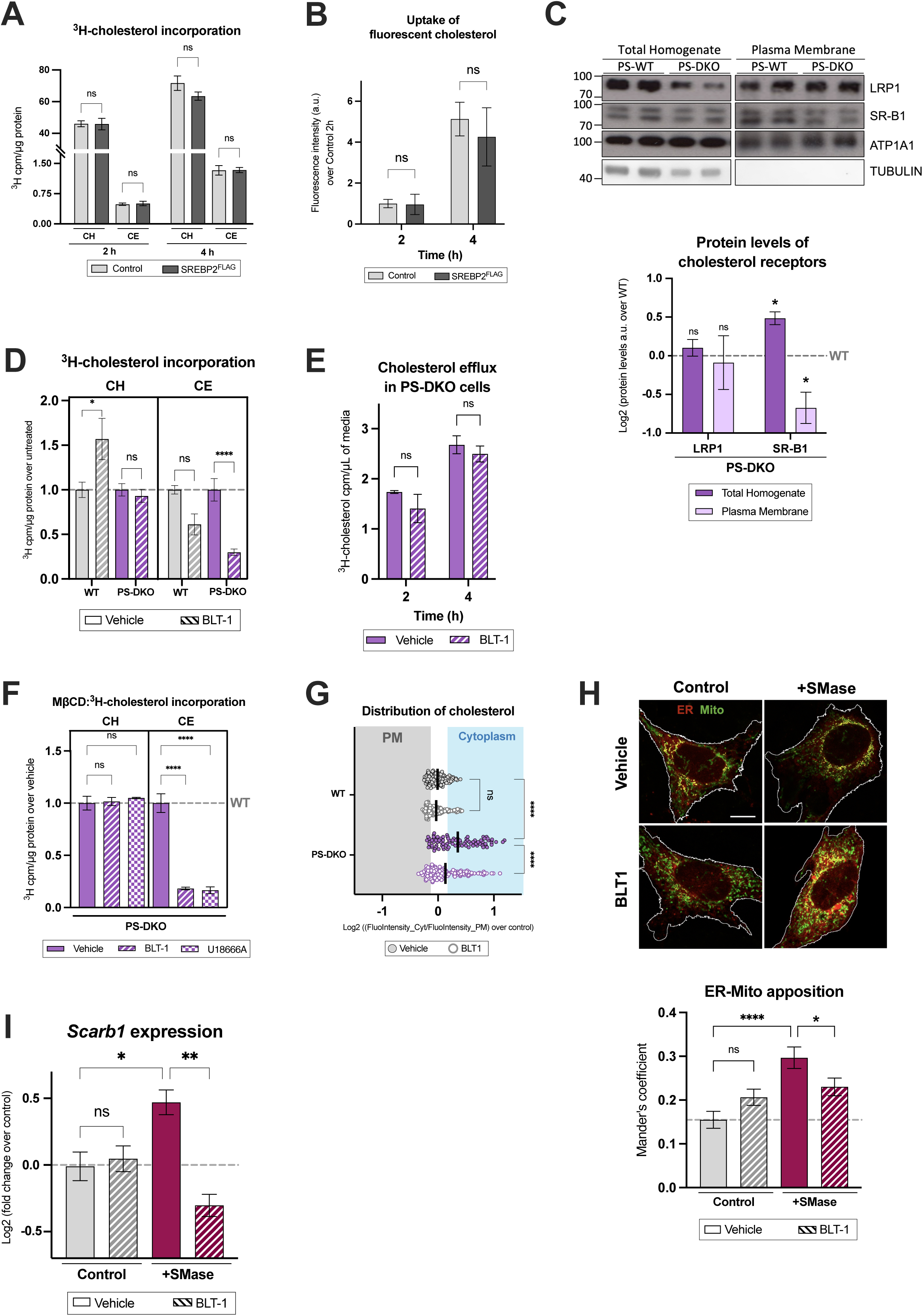
SR-B1-mediated uptake of cholesterol induces MAM formation (cont.). **A.** Quantification of exogenous ^3^H-cholesterol uptake (CH) and esterification (CE) in MEF cells overexpressing SREBP2-FLAG at the indicated times (*n = 3 biological replicates*). **(B)** Quantification of the internalization of fluorescent NBD-cholesterol in MEF cells overexpressing SREBP2-FLAG at the indicated times (*n = 3 biological replicates*). **(C)** Representative immunoblot of cholesterol receptors (LRP1, SR-B1) levels in PS-DKO cells. Sizes at left in kDa. Below, quantification of protein levels normalized to the loading control (TUBULIN for total homogenate; ATP1A1 for plasma membrane) over WT (*n = 3 biological replicates*). **(D)** Quantification of exogenous ^3^H-cholesterol uptake (CH) and esterification (CE) in PS-DKO cells treated with 10 μM BLT-1 for 16h (*n = 3 biological replicates*). **(E)** Quantification of cholesterol efflux in PS-DKO cells exposed to ^3^H-cholesterol for 3h, treated with 10 μM BLT-1 (*n = 3 biological replicates*). **(F)** Uptake of MβCD:^3^H-cholesterol complexes (CH) and cholesterol esterification (CE) in PS-DKO cells treated with 10 μM BLT-1 or 1 μM U18666A for 16h (*n = 3 biological replicates*). **(G)** Quantification of cholesterol distribution in PS-DKO cells visualized by filipin staining before and after treatment with 10 μM BLT-1 for 6h, (*n = 87-138 cells per condition*). **(H)** Representative image of ER (red)-mitochondria (Mito, green) apposition in MEF WT cells exposed to 1 U/mL SMase for 3h before and after treating with 10 μM BLT-1. Scale bar = 10 μm. Below, quantification of the co-localization coefficient (*n = 43 cells per condition*). **(I)** Measurement of mRNA expression levels of *Scarb1* in MEF WT cells exposed to 1 U/mL SMase for 3h, treated with 10 μM BLT-1 (*n = 3 biological replicates*). All data are expressed as the mean ± SEM. * p<0.05, ** p<0.01, **** p<0.0001.

**Fig S6.**
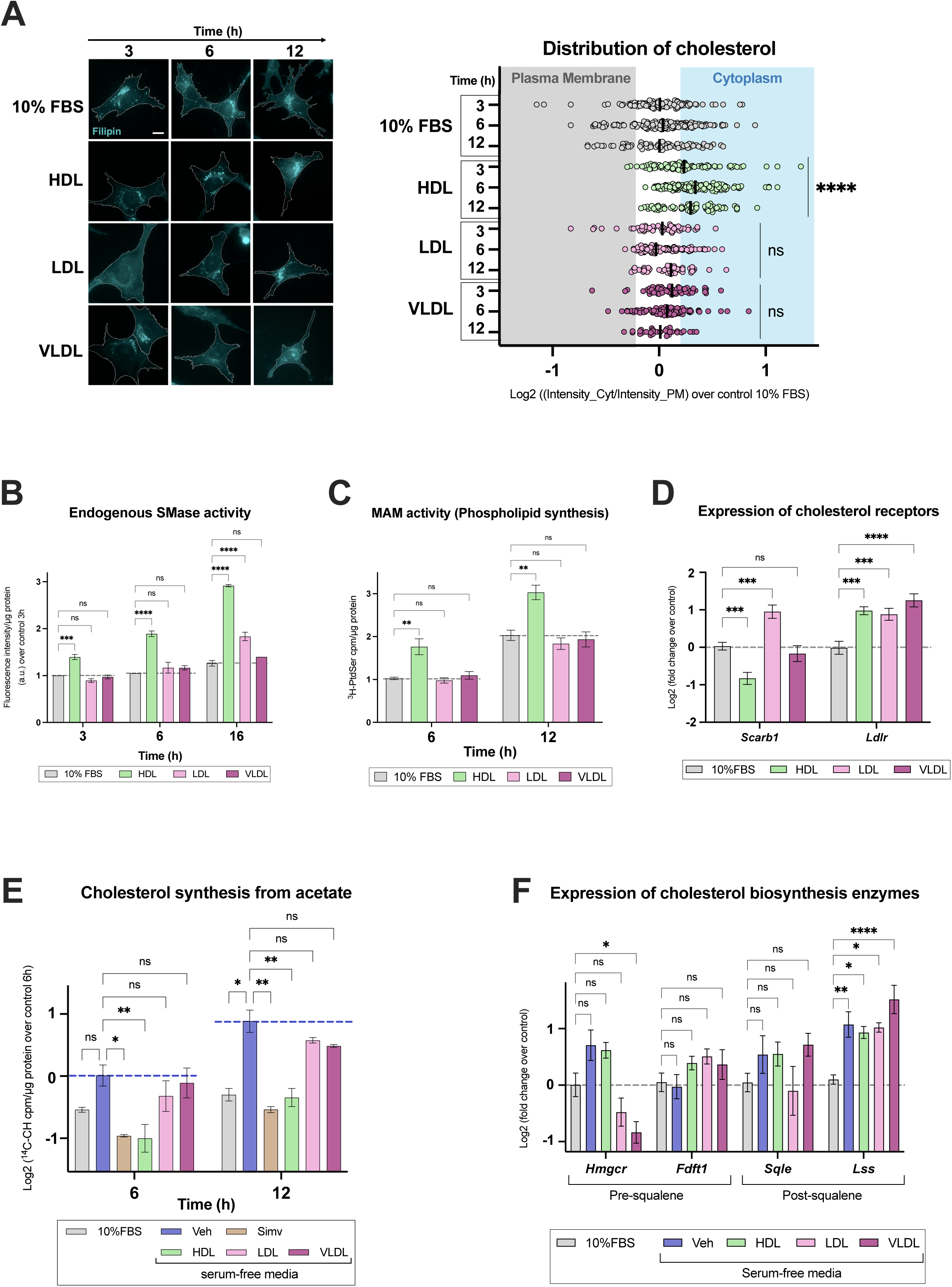
HDL–SR-B1 signaling induces MAM formation over time. All experiments were performed in MEF cells exposed to 5 ng/mL of different lipoproteins (HDL, LDL, VLDL) for 3, 6 or 12h. **(A)** Representative images of cholesterol distribution by filipin staining in cells under the indicated conditions, scale bar = 10 μm. Below, quantification of the distribution of cholesterol based on filipin staining over control 10% FBS (*n = 31-149 cells per condition*). (**B)** Measurement of endogenous SMase activity (*n = 3 biological replicates*). **(C)** Measurement of MAM activity, assessed by phosphatidylserine (PtdSer) synthesis (*n = 3 biological replicates*). **(D)** Measurement of mRNA expression levels of cholesterol receptors (*Scarb1* and *Ldlr*) (*n = 3 biological replicates*). **(E)** Measurement of *de novo* cholesterol synthesis from ^14^C-acetate in serum-free cells under the indicated conditions (*n = 3 biological replicates*). **(F)** Measurement of mRNA expression levels of cholesterol biosynthesis enzymes (*n = 9 biological replicates*). All data are expressed as the mean ± SEM. * p<0.05, ** p<0.01, *** p<0.001, **** p<0.0001.

## REFERENCES

1. Prinz, W.A., A. Toulmay, and T. Balla, The functional universe of membrane contact sites. Nat Rev Mol Cell Biol, 2020. 21(1): p. 7–24.

2. Bui, V., et al., Mitochondria-associated membranes (MAMs): molecular organization, cellular functions, and their role in health and disease. FEBS Open Bio, 2026. 16(1): p. 11–24.

3. Vance, J.E., Inter-organelle membrane contact sites: implications for lipid metabolism. Biol Direct, 2020. 15(1): p. 24.

4. Khodadadi, E., P. Chaturvedi, and M. Moradi, Comprehensive Insights into the Cholesterol-Mediated Modulation of Membrane Function Through Molecular Dynamics Simulations. Membranes (Basel), 2025. 15(6).

5. Isik, O.A. and O. Cizmecioglu, Rafting on the Plasma Membrane: Lipid Rafts in Signaling and Disease. Adv Exp Med Biol, 2023. 1436: p. 87–108.

6. Hoffmann, C., et al., Membrane-protein-mediated phase separation orchestrates organelle contact sites. Mol Cell, 2026. 86(1): p. 135–149.e9.

7. Sezgin, E., et al., The mystery of membrane organization: composition, regulation and roles of lipid rafts. Nat Rev Mol Cell Biol, 2017. 18(6): p. 361–374.

8. Doktorova, M., et al., Cell membranes sustain phospholipid imbalance via cholesterol asymmetry. Cell, 2025. 188(10): p. 2586–2602.e24.

9. Ikonen, E., Mechanisms of cellular cholesterol compartmentalization: recent insights. Curr Opin Cell Biol, 2018. 53: p. 77–83.

10. Hao, M., et al., Vesicular and non-vesicular sterol transport in living cells. The endocytic recycling compartment is a major sterol storage organelle. J Biol Chem, 2002. 277(1): p. 609–17.

11. Cui, D., et al., Cholesterol metabolism: molecular mechanisms, biological functions, diseases, and therapeutic targets. Mol Biomed, 2025. 6(1): p. 72.

12. Sharpe, L.J., H.W. Coates, and A.J. Brown, Post-translational control of the long and winding road to cholesterol. J Biol Chem, 2020. 295(51): p. 17549–17559.

13. Luo, J., et al., Routes and mechanisms of post-endosomal cholesterol trafficking: A story that never ends. Traffic, 2017. 18(4): p. 209–217.

14. Xiao, X., et al., Selective Aster inhibitors distinguish vesicular and nonvesicular sterol transport mechanisms. Proc Natl Acad Sci U S A, 2021. 118(2).

15. Zhu, P., L. Xiao, and Y. Wang, Mechanisms of high-density lipoprotein in regulating blood-brain barrier function: insights and implications. Fluids Barriers CNS, 2025. 22(1): p. 113.

16. Zhang, Q., Y. Ke, and H. Hong, HDL and Lipid Metabolism. Adv Exp Med Biol, 2022. 1377: p. 49–61.

17. Yu, H., HDL and Scavenger Receptor Class B Type I (SRBI). Adv Exp Med Biol, 2022. 1377: p. 79–93.

18. von Eckardstein, A., et al., High-density lipoprotein revisited: biological functions and clinical relevance. Eur Heart J, 2023. 44(16): p. 1394–1407.

19. Giacona, J.M., et al., HDL-C and apolipoprotein A-I are independently associated with skeletal muscle mitochondrial function in healthy humans. Am J Physiol Heart Circ Physiol, 2024. 326(4): p. H916–H922.

20. Montesinos, J., et al., The Alzheimer’s disease-associated C99 fragment of APP regulates cellular cholesterol trafficking. EMBO J, 2020. 39(20): p. e103791.

21. Haeffner, E.W. and U. Wittmann, Cholesterol-induced growth stimulation, macromolecule synthesis, and increased phosphoinositide metabolism of ascites tumour cells in culture. Cell Signal, 1999. 11(11): p. 821–9.

22. Harned, T.C., et al., Acute ACAT1/SOAT1 Blockade Increases MAM Cholesterol and Strengthens ER-Mitochondria Connectivity. Int J Mol Sci, 2023. 24(6).

23. Montesinos, J., E. Area-Gomez, and M. Schlame, Analysis of phospholipid synthesis in mitochondria. Methods Cell Biol, 2020. 155: p. 321–335.

24. Slotte, J.P. and E.L. Bierman, Depletion of plasma-membrane sphingomyelin rapidly alters the distribution of cholesterol between plasma membranes and intracellular cholesterol pools in cultured fibroblasts. Biochem J, 1988. 250(3): p. 653–8.

25. Zou, X., et al., SREBPs in Metabolic Reprogramming and Disease: Mechanisms and Therapeutic Potential. FASEB J, 2026. 40(12): p. e72037.

26. Area-Gomez, E., et al., Upregulated function of mitochondria-associated ER membranes in Alzheimer disease. EMBO J, 2012. 31(21): p. 4106–23.

27. Brown, M.S. and J.L. Goldstein, A proteolytic pathway that controls the cholesterol content of membranes, cells, and blood. Proc Natl Acad Sci U S A, 1999. 96(20): p. 11041–8.

28. Yu, M., et al., Contributions of a disulfide bond and a reduced cysteine side chain to the intrinsic activity of the high-density lipoprotein receptor SR-B1. Biochemistry, 2012. 51(50): p. 10044–55.

29. Frolov, A., et al., High density lipoprotein-mediated cholesterol uptake and targeting to lipid droplets in intact L-cell fibroblasts. A single- and multiphoton fluorescence approach. J Biol Chem, 2000. 275(17): p. 12769–80.

30. Voeltz, G.K., et al., Making the connection: How membrane contact sites have changed our view of organelle biology. Cell, 2024. 187(2): p. 257–270.

31. Elustondo, P., L.A. Martin, and B. Karten, Mitochondrial cholesterol import. Biochim Biophys Acta Mol Cell Biol Lipids, 2017. 1862(1): p. 90–101.

32. Doghman-Bouguerra, M. and E. Lalli, The ER-mitochondria couple: In life and death from steroidogenesis to tumorigenesis. Mol Cell Endocrinol, 2017. 441: p. 176–184.

33. Wall, C.T.J., et al., *Mitochondrial respiratory chain dysfunction alters ER sterol sensing and* mevalonate *pathway activity*. J Biol Chem, 2022. 298(3): p. 101652.

34. Infante, R.E. and A. Radhakrishnan, Continuous transport of a small fraction of plasma membrane cholesterol to endoplasmic reticulum regulates total cellular cholesterol. Elife, 2017. 6.

35. Endapally, S., et al., Molecular Discrimination between Two Conformations of Sphingomyelin in Plasma Membranes. Cell, 2019. 176(5): p. 1040–1053.e17.

36. Lange, Y. and T.L. Steck, Quantitation of the pool of cholesterol associated with acyl-CoA:cholesterol acyltransferase in human fibroblasts. J Biol Chem, 1997. 272(20): p. 13103–8.

37. Naito, T., et al., Movement of accessible plasma membrane cholesterol by the GRAMD1 lipid transfer protein complex. Elife, 2019. 8.

38. Sala-Vila, A., et al., Interplay between hepatic mitochondria-associated membranes, lipid metabolism and caveolin-1 in mice. Sci Rep, 2016. 6: p. 27351.

39. Ferrari, A. and P. Tontonoz, Nonvesicular cholesterol transport in physiology. J Clin Invest, 2025. 135(6).

40. Baumann, N.A., et al., Transport of newly synthesized sterol to the sterol-enriched plasma membrane occurs via nonvesicular equilibration. Biochemistry, 2005. 44(15): p. 5816–26.

41. Webb, N.R., et al., The fate of HDL particles in vivo after SR-B1-mediated selective lipid uptake. J Lipid Res, 2002. 43(11): p. 1890–8.

42. Naito, T., et al., Regulation of cellular cholesterol distribution via non-vesicular lipid transport at ER-Golgi contact sites. Nat Commun, 2023. 14(1): p. 5867.

43. Kristiana, I., H. Yang, and A.J. Brown, Different kinetics of cholesterol delivery to components of the cholesterol homeostatic machinery: implications for cholesterol trafficking to the endoplasmic reticulum. Biochim Biophys Acta, 2008. 1781(11-12): p. 724–30.

44. Montesinos, J. and E. Area-Gomez, Isolation of mitochondria-associated ER membranes. Methods Cell Biol, 2020. 155: p. 33–44.

45. Pera, M., et al., Increased localization of APP-C99 in mitochondria-associated ER membranes causes mitochondrial dysfunction in Alzheimer disease. EMBO J, 2017. 36(22): p. 3356–3371.

